# TENT5C extends *Odf1* poly(A) tail to sustain sperm morphogenesis and fertility

**DOI:** 10.1101/2025.03.20.644152

**Authors:** Marine Baptissart, Ankit Gupta, Alexander C. Poirot, Brian N. Papas, Marcos Morgan

## Abstract

Changes in the poly(A) tail length of *Odf1* and other transcripts critical for male fertility have been linked to translational activation during sperm formation ^1–3^. The mRNA poly(A) polymerase TENT5C is required for fastening the flagellum to the sperm head, but its role in shaping the poly(A) tail profile of the spermatid transcriptome remains limited ^4,5^. Here, we comprehensively document how changes in mRNA poly(A) tail length across the transcriptome reflect transcript metabolism in spermatids. In the absence of TENT5C polymerase activity, the poly(A) tail length of *Odf1* transcripts is reduced, and the local distribution of ODF1 proteins in spermatids is disrupted. We show that mice expressing a catalytically inactive TENT5C produce headless spermatozoa with outer dense fibers detached from the axoneme, and other flagellar abnormalities associated with ODF1 deficiency ^6^. We propose that TENT5C poly(A) polymerase activity regulates the spatial translation of *Odf1* mRNAs during spermiogenesis, a process critical for sperm morphogenesis and fertility. These findings highlight the power of poly(A) tail profiling to identify abnormal mRNA processing causative of infertility.

## MAIN

Spermiogenesis is the final step of spermatogenesis, where haploid spermatids exchange histones for protamines and elongate to produce sperm with a condensed nucleus and a flagellum. Spermiogenesis is one of the rare events where post-transcriptional regulation occurs in the absence of transcription ^7^. Transcription ceases in stage 10 round spermatids (RS) and remains silent for the rest of the differentiation process ^8,9^. In this context, the mRNAs required to orchestrate the late differentiation program are synthesized early on and stabilized before translational activation ^1–3,10–12^. However, the post-transcriptional mechanisms controlling the temporal disconnect between mRNA transcription and translation during spermiogenesis are poorly understood. The poly(A) tail is a stretch of A nucleotides (nts) added co-transcriptionally to the 3’-end of most eukaryotic mRNAs ^13^. Once in the cytoplasm, poly(A)-binding proteins (PABPs) coat the poly(A) tail to favor transcript stability and translation ^14^. Deadenylase complexes progressively peel off PABPs, leading to poly(A) tail shortening and mRNA decay ^15^. A correlation between poly(A) tail length and mRNA stability and translation has been clearly established in cells where transcription is silenced ^3,16–19^. The most well-characterized example is the elongation of poly(A) tails during oocyte activation, which leads to translational activation ^20–23^. Early reports have shown that during the transition from pachytene spermatocytes to RS, the poly(A) tail of transcripts critical for spermiogenesis is extended by up to 100 additional nts, a process proposed to favor their stability ^10–12^. As the RS differentiate into elongated spermatids (ES), mRNAs encoding for key structural and chromatin factors show poly(A) tails shortening from ∼150- to ∼30-nt long, a dynamic associated with their translational activation ^1–3^. A recent study examined transcriptome-wide poly(A) tail dynamics during spermiogenesis using long-read sequencing ^24^. However, unlike previous reports, transcripts were proposed to be stored and translationally repressed with tails ∼30-nt long. Thus, how the global poly(A) tail dynamics regulate mRNA metabolism and spermatid differentiation is still unclear. In recent years, several terminal nucleotidyltransferases (TENTs) have been shown to be required to sustain gametogenesis ^4,25–27^. Among them, the poly(A) polymerase TENT5C, highly abundant in the manchette of ES, was shown to be required for male fertility ^4^. *Tent5c null* males produce headless spermatozoa due to defective spermiation, the process by which the ES are released from the seminiferous tubules. This phenotype was not associated with global changes in mRNA accumulation when sampling whole testis, suggesting that the poly(A) polymerase activity of TENT5C might not be required for spermatogenesis ^4^. More recently, a poly(A) profile of whole testes from *Tent5c null* animals has shown changes in poly(A) tail length for several transcripts expressed in various testicular cell types ^5^. Thus, the role of TENT5C poly(A) polymerase, if any, in late spermatid differentiation where the phenotype is observed remains limited.

In this study, we characterize transcriptome-wide poly(A) tail dynamics during spermiogenesis and its impact on mRNA metabolism and fertility. After meiosis, most of the RS transcriptome is stabilized with tails ∼150-nt long. As the spermatids elongate, the translational activity of mRNAs is reflected by the accumulation of transcripts with ∼60-nt long poly(A) tails. Additionally, we show that the poly(A) polymerase activity of TENT5C is required for fertility and spermiogenesis; male mice expressing catalytically dormant TENT5C (*Tent5c^dcat/dcat^*) are sterile and produce headless spermatozoa with abnormal flagellar morphologies. We identified outer dense fiber of sperm tails 1 (*Odf1)* as one of the few transcripts dependent on the poly(A) polymerase activity of TENT5C in ES. In *Tent5c^dcat/dcat^* mice, *Odf1* showed a reduction in poly(A) tail length with an increased proportion of transcripts with tails ∼60-nt long. This profile is associated with the permissive accumulation of ODF1 protein across the ES cytoplasm. The aberrant localization of ODF1 can causally explain the bent and fragmented flagella observed in the infertile *Tent5c^dcat/dcat^* mice. These results show that poly(A) profiling of germ cells can be used to identify specific transcripts that when dysregulated cause male infertility.

### The characteristic accumulation of ∼150-nt long poly(A) tails in spermatids is associated with the stabilization of the transcriptome

Seminal studies have shown that transcripts required for sperm metabolism, nuclear compaction and flagella formation show dynamic poly(A) tail length profiles during spermiogenesis ^1–3,10–12^. To understand whether this behavior is characteristic of a few transcripts or prevalent to the spermatid transcriptomes, we used a well-established FACS sorting strategy to isolate germ cell populations enriched with pachytene and diplotene spermatocytes (P/D spermatocytes), RS or ES ^28^(**Extended Data Fig. 1a**). The purity of isolated populations was quantitatively determined by microscopy using fluorescent markers (**Extended Data Fig. 1a to c**). Late spermatocytes were identified with the meiotic marker synaptonemal complex protein 3 (SYCP), while acrosomal structures characteristic of RS and ES were stained with Peanut agglutinin (PNA). DAPI was used to stain DNA. About 50% of the cells from the sorting gate II were P/D spermatocytes, while the gates IIIa and IIIb were enriched with more than 80% of RS and ES, respectively (**Extended Data Fig. 1b and c**). Cell identity was further confirmed through direct RNA sequencing. Principal component analysis showed clustering of the populations according to their expected cell types (**Extended Data Fig. 1d**). We next mapped each sequenced transcriptome to cell clusters from a whole testis single-cell RNA sequencing dataset to computationally predict the cell type composition ^29^. Each individual sample displayed a strong signature corresponding to the anticipated germ cell transcriptome (**Extended Data Fig. 1e**).

To characterize the global poly(A) profile of each germ cell population, we quantified poly(A) tail length from our direct RNA-seq datasets using Nanopolish ^30^. The transcriptome of P/D spermatocytes had predominantly poly(A) tails ∼60-nt long, while the transcriptome of RS peaked with poly(A) tails ∼150-nt long (**Fig. 1a**). The transcripts encoding for lactate dehydrogenase C (*Ldhc*) and ornithine decarboxylase 1 (*Odc1*) showed poly(A) tails longer in RS than in P/D spermatocytes as previously reported ^10,11^(**Fig. 1b, Extended Data Fig. 1f**). Our global approach revealed that the transition to long poly(A) tail profiles was not limited to few targets; the mean poly(A) tail length significantly increased in more than 64% (2469/3825) of the captured transcripts during the transition from P/D spermatocytes to RS (**Fig. 1b**). Only ∼1% (54/3825) of the mRNAs showed a significant reduction in poly(A) tail length from P/D to RS, with the mRNA of cyclin A1 (*Ccna1*), a key regulator of meiotic progression, showing a pronounced shortening ^31^ (**Fig. 1b, Extended Data Fig. 1f**). Thus, the transition from spermatocytes to spermatids is characterized by the global lengthening of transcript poly(A) tails from ∼60 to ∼150 nts. Since active transcription ceases early during spermatid elongation ^8,9^, the long poly(A) tail profile of RS has been proposed to provide long-term stability to transcripts so they can be used days later to support ES differentiation. Given the transcriptional silencing in late spermiogenesis, we used differential RNA accumulation between ES and RS to infer RNA stability during spermiogenesis; Downregulated transcripts were considered unstable, while upregulated transcripts were deemed stable (**Extended Data Fig.1g and Extended Data Table 1**). Poly(A) tail profiling of RS revealed a higher accumulation of ∼150-nt long tails for stable transcripts compared to the rest of the transcriptome (**Fig. 1c, Extended Data Fig. 1h**). This observation was confirmed at the single transcript level. Changes in mRNA accumulation across the transcriptome positively correlated with the proportion of ∼150-nt long poly(A) tails in RS (**Fig. 1d**). The transcripts encoding for SMCP, PRM2 and ODF1, key factors during late spermiogenesis, were among those with highest stability and accumulation of ∼150-nt long tails in RS (**Fig. 1d**). Conversely, the unstable *Ccna1* transcripts showed a low proportion of ∼150-nt tails in RS (**Fig. 1d**). When grouping individual transcripts according to their stability, the proportion of mRNAs with ∼150-nt long poly(A) tails was in average statistically higher for stable transcripts, while unstable transcripts showed less ∼150-nt tails compared to the rest of the transcriptome (**Fig. 1e**). Together, these results show that the accumulation of poly(A) tails ∼150-nt long in RS predicts global transcript stability through spermiogenesis.

**Figure 1.**
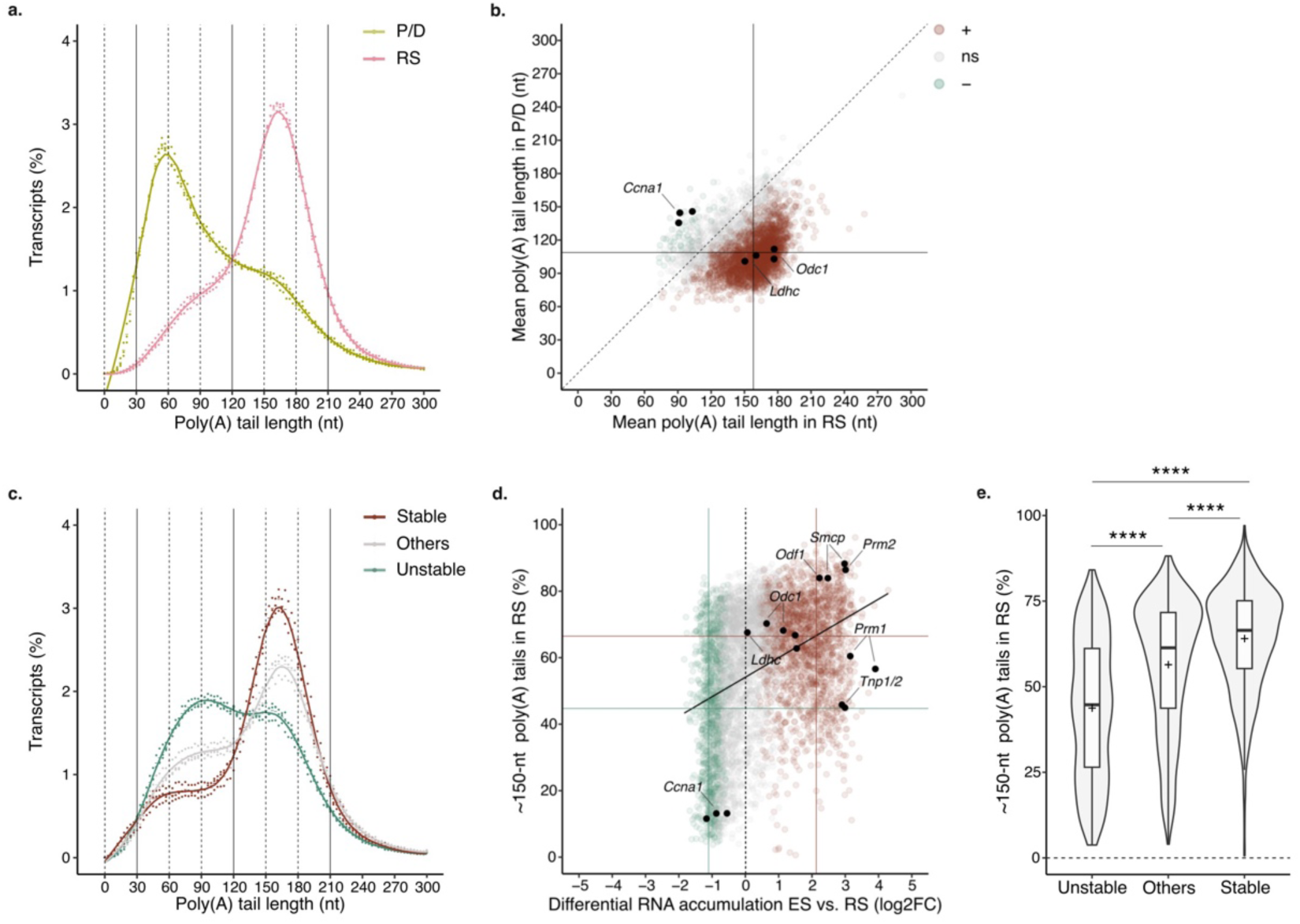
The characteristic accumulation of ∼150-nt long poly(A) tails in spermatids is associated with the stabilization of the transcriptome. **(a)** Poly(A) tail length density plot of pachytene/diplotene spermatocyte (P/D, green) and round spermatid (RS, pink) transcriptomes. Dots indicate values for individual biological replicates. The bars indicate the relative mean percentage of transcripts for each poly(A) tail length. The local polynomial regression fitting is shown as a solid line for each condition. nt: nucleotide. **(b)** Scatter plot comparing the mean poly(A) tail length of each transcript in pachytene/diplotene spermatocytes (P/D) and round spermatids (RS). Each dot represents an individual transcript. Transcripts significantly increasing (+) or decreasing (-) in poly(A) tail length during the transition are shown in brown or green, respectively; Student’s t-test, two-tailed, significance threshold p < 0.05. ns: non significant. *Odc1*, *Ldhc and Ccna1* transcripts are indicated in black. **(c)** Poly(A) tail length density plot as in (a) for stable, others, and unstable transcripts in round spermatids (RS) shown in brown, grey and green, respectively. Transcript stability through spermiogenesis is defined by the differential mRNA accumulation between elongated spermatids (ES) and round spermatids (RS). **(d)** Scatter plot comparing fold change in transcript differential accumulation between round spermatids (RS) and elongated spermatids (ES) to the percentage of reads per transcript with poly(A) tail ∼150-nt long in RS. Each dot represents an individual transcript. Upregulated (up), others, and downregulated transcripts (down) are indicated in brown, grey and green, respectively. Solid lines indicate the median of both variables for upregulated and downregulated mRNAs in brown and green, respectively. The linear fit is shown in black. Specific transcripts are indicated in black. **(e)** Violin plot showing the proportion of reads per transcript with poly(A) tails ∼150-nt long in round spermatids (RS). Transcripts are grouped by stability. The width of violins show the density of individual transcripts. The overlaid box plots display means as crosses and medians as lines; the boxes indicate the first and third quartiles and the bars indicate the 10^th^ and 90^th^ percentiles. Krustal-Wallis test using Bonferroni correction for mutilple testing. ****p < 0.0001. n=3 biological replicates per condition.

### Changes in poly(A) profiles reflect the degree of mRNA translational activation during spermiogenesis

Previous works have shown that the length of *Tnp1*, *Tnp2*, *Prm1*, *Prm2*, *Smcp* and *Odf1* poly(A) tails decreases during spermiogenesis progression from RS to ES ^1–3^. To determine the extent of poly(A) shortening, we compared the poly(A) profiles of RS and ES transcriptomes. Although the transcriptome of ES retained a prominent ∼150-nt long poly(A) tail peak, we observed a significant increase in the frequency of transcripts with ∼60-nt poly(A) tails compared to RS (**Fig. 2a**). The increase was mostly driven by stable transcripts which showed a ∼1-fold increase in the proportion of ∼60-nt long poly(A) tails from RS to ES, while unstable transcripts only showed a modest increase (**Extended Data Fig. 2a to c**). When analyzed individually, more than 44% (2362/5290) of captured transcripts showed a significant reduction in mean poly(A) tail length during spermiogenesis progression (**Fig. 2b**). Consistent with previous findings, the poly(A) tail length of the transcripts encoding for protamines, transition proteins, ODF1 and SMCP were significantly reduced in ES compared to RS ^1–3^(**Fig. 2b**). However, the amplitude of their tail shortening was modest compared to most transcripts, since although the fraction of ∼60-nt long tails increased significantly, most transcripts were retained with ∼150-nt long tails (**Fig. 2b and c**). In contrast, *Ldhc* and *Odc1*, which largely accumulated with ∼150-nt long tails in RS, showed a pronounced poly(A) tail shortening back to ∼60 nts in ES (**Fig. 2b and Extended Data Fig. 2d**). In sum, although most of the transcript poly(A) tails are shortened during spermiogenesis, the amplitude of the shortening varies widely between transcripts.

**Figure 2.**
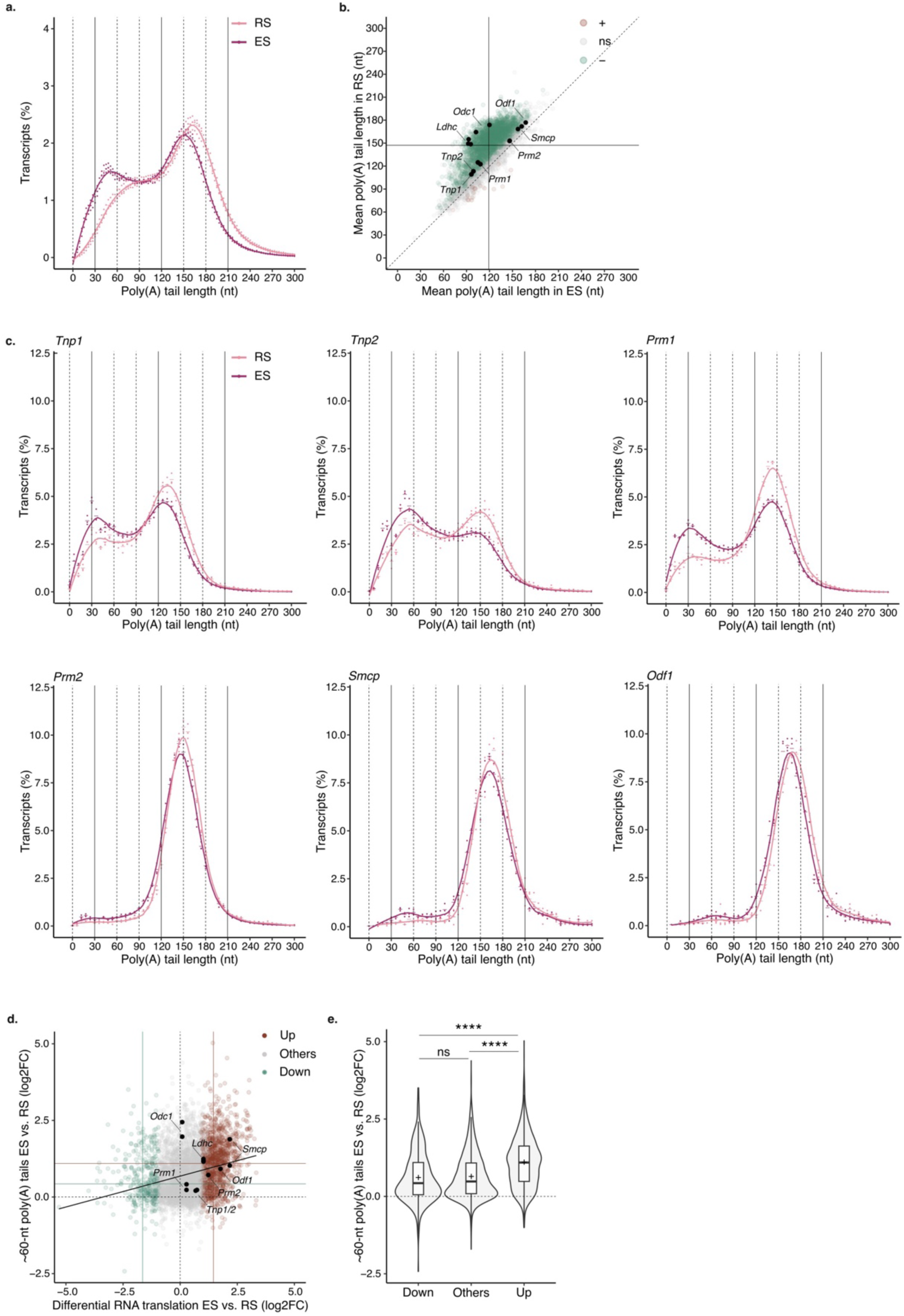
Changes in poly(A) profiles reflect the degree of mRNA translational activation during spermiogenesis. **(a)** Poly(A) tail length density plot for round spermatid (RS, pink) and elongated spermatid (ES, magenta) transcriptomes. Dots indicate values for individual biological replicates. The bars indicate the relative mean percentage of transcripts for each poly(A) tail length. The local polynomial regression fitting is shown as a solid line for each condition. nt: nucleotide. **(b)** Scatter plot comparing the mean poly(A) tail length of each transcript in round spermatids (RS) and elongated spermatids (ES). Each dot represents an individual transcript. Transcripts significantly increasing (+) or decreasing (-) in poly(A) tail length during the transition are shown in brown or green, respectively; Student’s t-test, two-tailed, significance threshold p < 0.05. ns: non significant. Specific transcripts are indicated in black. **(c)** Poly(A) tail length density plots as in (a) for *Tnp1*, *Tnp2*, *Prm1*, *Prm2*, *Smcp*, and *Odf1*. **(d)** Scatter plot comparing fold change in transcript differential translation to the fold change in the percentage of reads per transcript with poly(A) tail ∼60-nt long between RS and ES. Each dot represents an individual transcript. Transcript with increased (Up), unchanged (Others) or decreased (Down) levels of translation are indicated in brown, grey and green, respectively. Solid lines indicate the median of both variables for upregulated and downregulated mRNAs in brown and green, respectively. The linear fit is shown in black. Specific transcripts are indicated in black. **(e)** Violin plot showing the fold change in the percentage of reads per transcript with poly(A) tail ∼60-nt long between round spermatids (RS) and elongated spermatids (ES). Transcripts are grouped by differential translation. The width of violins show the density of individual transcripts. The overlaid box plots display means as crosses and medians as lines; the boxes indicate the first and third quartiles and the bars indicate the 10^th^ and 90^th^ percentiles. Krustal-Wallis test using Bonferroni correction for mutilple testing. ****p < 0.0001. n=3 biological replicates per condition.

The shortening of the poly(A) tails during spermiogenesis has been previously associated with the translational activation of *Tnp1*, *Tnp2*, *Prm1*, *Prm2*, *Smcp* and *Odf1* ^1–3^. To understand whether the association is true across the transcriptome, we used previously published data to integrate translational activation between RS and ES to our global poly(A) tail length analysis ^32^. Transcripts accumulating with ∼60-nt long tails during spermiogenesis progression were also the ones showing significant translational activation compared to the rest of the transcriptome (**Fig. 2d and e, Extended Data Fig. 2e**). Among the transcripts with poly(A) tail shortening to ∼60 nts, *Prm2*, *Smcp* and *Odf1* showed a ∼2 fold increase in translational activation (**Fig. 2d**). A closer inspection to their individual poly(A) profiles revealed that despite their transition to ∼60 nts, most transcripts are retained with tails ∼150-nt long (**Fig. 2c**). In contrast, transition proteins and *Prm1* transcripts showed a lower abundance of ∼150-nt long tails, and modest translational activation during spermatid elongation (**Fig. 2c and d**). This indicates that while the transition to ∼60-nt poly(A) tail is a signature of translational acceleration, the amplitude of the translational activation increases with the proportion of transcripts retained with ∼150-nt tails. Taken together, these results support a model in which the most stable transcripts are stored with tails 150-nt long in RS. During spermatids elongation, these transcripts become translationally active with tails ∼150-nt long. Their poly(A) tail is gradually shortened during translation leading to accumulation of ∼60-nt tails before transcripts get processed for further deadenylation and decay.

### TENT5C catalytic activity is required for spermiogenesis

We next sought to identify the enzymes responsible for poly(A) tail elongation during spermiogenesis. TENT5C is a non-canonical poly(A) polymerase shown to be highly expressed in ES ^4^. We confirmed TENT5C localization using flow cytometry by tracking the fluorescent signal of germ cells from mice expressing a *Tent5c* knock-in GFP reporter allele ^33^. None of the gates enriched in early or late meiotic cells showed differential fluorescence between *Tent5c^gfp/gfp^* and wild-type mice, indicating the absence of significant TENT5C accumulation during meiosis (**Extended Data Fig. 3a and b**). However, TENT5C fluorescence was detected in isolated RS, with levels significantly increasing in the fraction enriched for ES (**Extended Data Fig. 3a and b**).

The presence of TENT5C is essential to complete spermiogenesis; *Tent5c^null/null^* mice produce headless spermatozoa and are sterile ^4,5^ (**Fig. 3a to c, Extended Data Fig. 3c to d**). Given the minor differences in RNA accumulation between whole testis of *Tent5c^null/null^* and wild-type mice, it has been proposed that the poly(A) polymerase activity of TENT5C may not be required to support spermatogenesis ^4^. Recently, an independent study reported that the sterility of *Tent5c^null/null^* mice is associated with minor poly(A) tail changes limited to a few transcripts when considering whole testis as a starting material for sequencing ^5^. To definitely answer whether the poly(A) polymerase activity of TENT5C is required for fertility, we took advantage of a previously characterized mouse strain expressing a catalytically dead version of TENT5C (*Tent5c^dcat/dcat^*) ^33^. The body mass of the *Tent5c^dcat/dcat^* and *Tent5c^null/null^* mice were comparable to wild-type (**Extended Data Fig. 3e**). Neither *Tent5c^dcat/dcat^* nor *Tent5c^null/null^* mice showed differences in reproductive organ appearance and weight (**Extended Data Fig. 3f**). However, *Tent5c^dcat/dcat^* mice, like the *Tent5c^null/null^* mice, were completely infertile (**Fig. 3a**). To understand the origin of the sterility, we inferred germ cell production from the total number of flagella isolated from the cauda epididymis. While germ cell production was not compromised in *Tent5c* mutants, significant morphological sperm defects were observed (**Fig. 3b and c**). Only headless spermatozoa were found in caudal epididymis extracts from *Tent5c^dcat/dcat^* and *Tent5c^null/null^* animals (**Fig. 3c**). Isolated flagella were identified with various degree of morphological abnormalities (**Fig. 3c and Extended Data Fig. 3g**). About 10% were bent with an acute angle often found between the midpiece and the principal piece; 50% of the flagella formed complete hairpins overlapping the mid and principal piece together and 40% displayed enlarged or coiled midpiece (**Fig. 3c**). Since histological analyses confirmed the presence of headless flagella in the ducts of the caput and cauda epididymis, we speculated that the morphological defect of *Tent5c^dcat/dcat^* sperm originated at the testis as previously described for *Tent5c^null/null^* sperm ^4^ (**Fig. 3d**). Like *Tent5c^null/null^* mice, *Tent5c^dcat/dcat^* males showed retention of sperm heads close to the basement membrane of stage VII-VIII seminiferous tubules (**Fig. 3e, Extended Data Fig. 3h**). In addition, a TUNEL assay marked apoptotic events in the basement membrane of stage VII-VIII tubules specifically for both *Tent5c^null/null^* and *Tent5c^dcat/dcat^* mutants (**Fig. 3f, Extended Data Fig. 3i**). In sum, *Tent5c^dcat/dcat^* males phenocopy the *Tent5c^null/null^* mice showing sterility associated with production of headless spermatozoa, demonstrating that the poly(A) polymerase activity of TENT5C is required for spermiogenesis and fertility.

**Figure 3.**
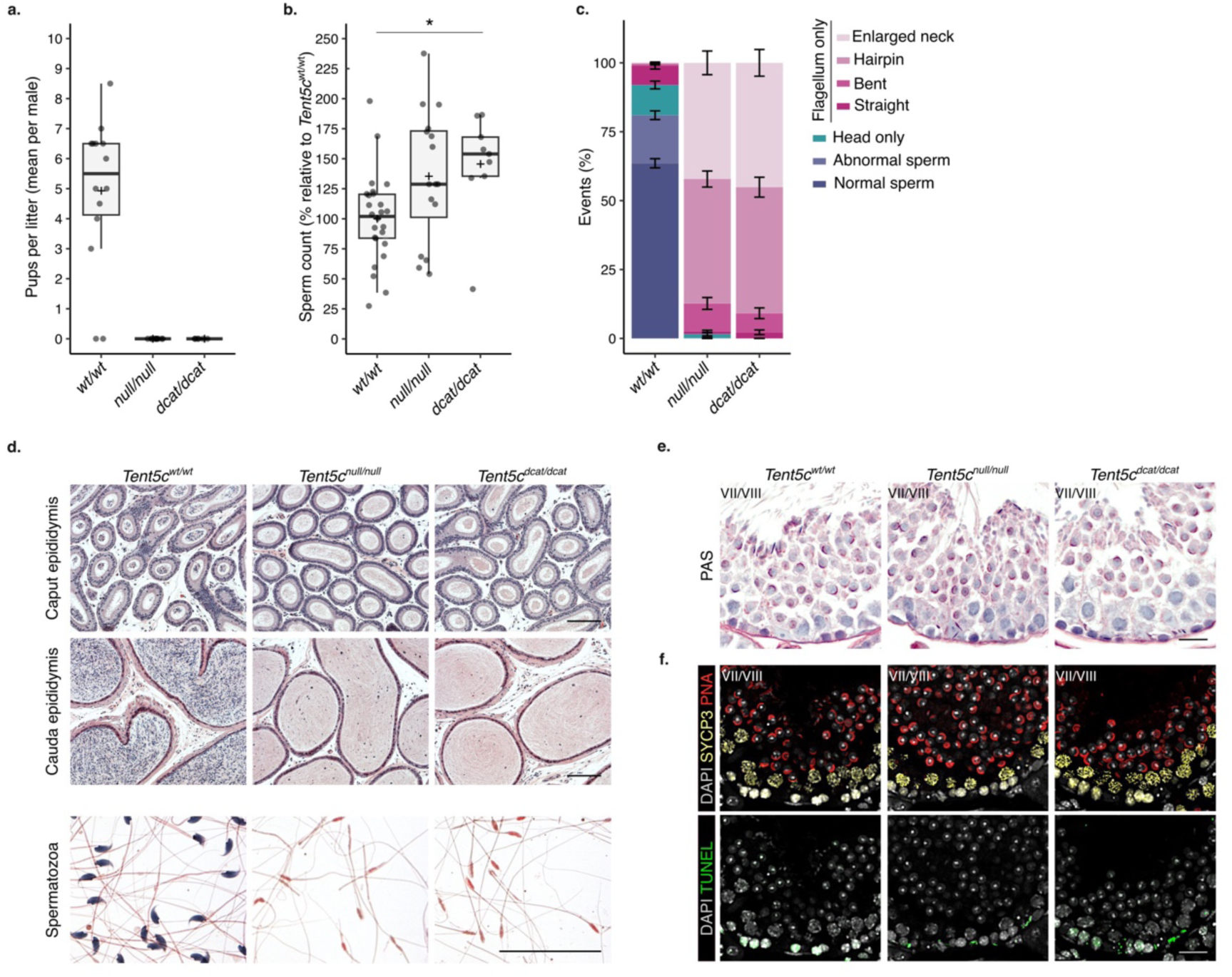
TENT5C poly(A) polymerase activity is required for fertility and spermiogenesis. **(a)** Box plot showing the average pups per litter obtained per adult male mice after mating with 2 control females. *Tent5c^null/null^* and *Tent5c^dcat/dcat^* mice are compared to *Tent5c^wt/wt^*. Each dot represents one mouse. The crosses display the means; the lines show the medians; the boxes indicate the first and third quartiles and the bars indicate the 10^th^ and 90^th^ percentiles. n=6-14 biological replicates per condition. **(b)** Box plot as in (a) showing the number of flagella from cauda epididymis for *Tent5c^null/null^* and *Tent5c^dcat/dcat^* mice expressed relative to *Tent5c^wt/wt^*. Krustal-Wallis test using Bonferroni correction for mutilple testing. *p<0.05. n=9-16 biological replicates per condition. **(c)** Bar plot showing the proportion of germ cell morphologies in semen isolated from the cauda epididymis of *Tent5c^wt/wt^, Tent5c^null/null^* and *Tent5c^dcat/dcat^*. Quantification from Hematoxylin and Eosin (H&E) staining using the classification presented in **Extended Data Fig. 3g**. n=8-18 biological replicates per condition. **(d)** Representative micrographs of epididymis cross sections (caput, upper; cauda, middle) and sperm from caudal epididymis (lower) stained with H&E. Sections from *Tent5c^null/null^* and *Tent5c^dcat/dcat^* mice are compared to *Tent5c^wt/wt^*. Sperm heads (nuclei) show a blue staining and flagella shades of pink. Scale, 40 μm. n=3 biological replicates per condition. **(e)** Representative micrographs as in (d) showing stage VII/VIII tubule cross sections stained with Periodic Acid Schiff (PAS). Arrows indicate sperm heads. Scale, 20 μm. n=3 biological replicates per condition. **(f)** Representative micrographs of stage VII/VIII tubule cross sections immunostaining from *Tent5c^wt/wt^*, *Tent5c^null/null^* and *Tent5c^dcat/dcat^*. SYCP immunostaining (yellow) marks spermatocytes; PNA labeling (red) marks spermatid acrosomes, TUNEL immunostaining (green) marks apoptotic cells and DAPI labeling (grey) marks nuclei. Arrows indicate apoptotic sperm heads engulfed in Sertoli cells. Scale, 20 μm. n=3 biological replicates per condition.

### *Insl3* and *Ptgds* poly(A) tail length depend on TENT5C activity in both elongated spermatids and Leydig cells

To identify the transcripts targeted by the poly(A) polymerase activity of TENT5C during spermiogenesis, we performed poly(A) profiling of RS and ES from wild-type and *Tent5c^dcat/dcat^* mice. In the absence of TENT5C poly(A) polymerase activity, we anticipated the poly(A) tail shortening of TENT5C targeted transcripts. The global poly(A) distributions of the RS and ES transcriptomes were almost identical between *Tent5c^dcat/dcat^* and wild-type mice (**Fig. 4a and b, and Extended Data Fig. 4a and b**). Moreover, no changes in transcript accumulation were observed between the two genotypes (**Extended Data Table 1**). These results show that TENT5C poly(A) polymerase activity is not required to shape the overall RNA poly(A) tail length profiles, nor transcript composition during spermiogenesis. Instead, we observed a significant poly(A) tail shortening restricted to a specific set of transcripts in RS and ES of *Tent5c^dcat/dcat^* mice (**Fig. 4b and Extended Data Fig. 4b**). Among them, insulin-like 3 (*Insl3*) and prostaglandin D2 synthase (*Ptgds*) mRNAs stand out as putative TENT5C targets. *Insl3* transcripts showed the highest reduction in mean poly(A) tail length in both RS and ES of *Tent5c^dcat/dcat^* mice compared to wild-type (**Fig. 4b and Extended Data Fig. 4b**). In ES, *Ptgds* transcript was the second transcript with the highest amplitude of poly(A) tail shortening (**Fig. 4b**). To better understand the nature of *Insl3* and *Ptgds* tail shortening, we examined their individual poly(A) length profiles. While *Insl3* and *Ptgds* mRNAs accumulate with tails ∼150-nt long in wild-type ES, the vast majority of *Insl3* and *Ptgds* transcripts shifted to ∼60-nt long poly(A) tails in *Tent5c^dcat/dcat^* mutants (**Fig. 4c**). Given that the accumulation of tails ∼60-nt long is associated with translational activation, we anticipated an increase in INSL3 protein abundance in *Tent5c^dcat/dcat^* ES. However, no ectopic signal for INSL3 protein was detected in any of the germ cells from wild-type and *Tent5c^dcat/dcat^* testes (**Fig. 4d**). Instead, INSL3 expression was restricted to Leydig cells as previously described ^5^. Thus, although TENT5C catalytic activity is required to sustain *Insl3* poly(A) tail length, the shortening of *Insl3* poly(A) tail in *Tent5c^dcat/dcat^* mutants appears not to be sufficient to trigger translation of *Insl3* transcript in ES.

**Figure 4.**
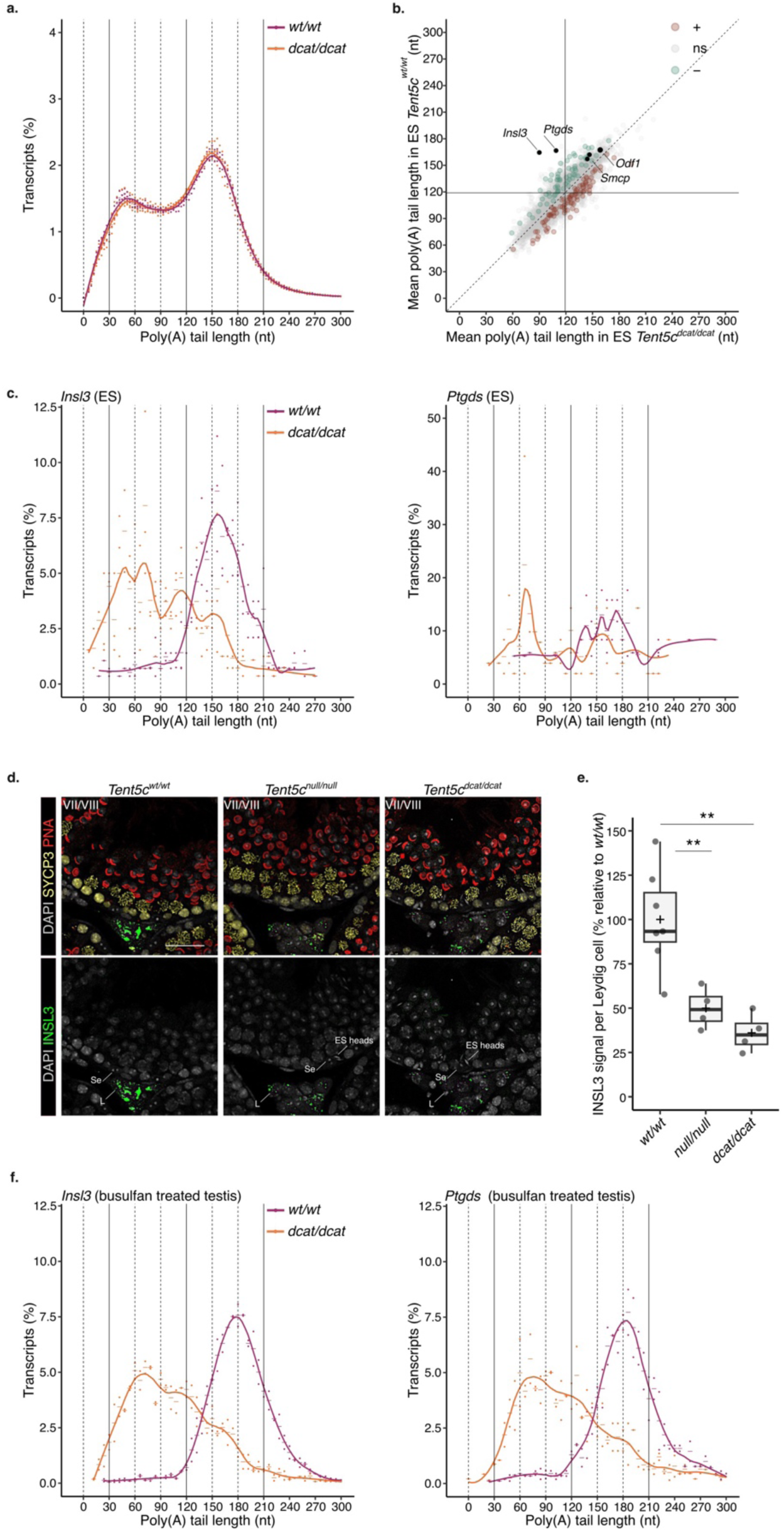
*Insl3* and *Ptgds* poly(A) tail length depend on TENT5C activity in both elongated spermatids and Leydig cells. **(a)** Poly(A) tail length density plot of the elongated spermatid (ES) transcriptomes of *Tent5c^wt/wt^* (magenta) *and Tent5c^dcat/dcat^* (orange) mice. Dots indicate values for individual biological replicates. The bars indicate the relative mean percentage of transcripts for each poly(A) tail length. The local polynomial regression fitting is shown as a solid line for each condition. nt: nucleotide. n=3 biological replicates per condition. **(b)** Scatter plot comparing the mean poly(A) tail length of each transcript in elongated spermatid (ES) of *Tent5c^wt/wt^* and *Tent5c^dcat/dcat^* mice. Each dot represents an individual transcript. Transcripts significantly increasing (+) or decreasing (-) in poly(A) tail length between genotypes are shown in brown or green, respectively; Student’s t-test, two-tailed, significance threshold p < 0.05. ns: non significant. n=3 biological replicates per condition. **(c)** Poly(A) tail length density plot as in (a) for *Insl3 and Ptgds* transcripts from the elongated spermatid (ES) of *Tent5c^wt/wt^* (magenta) *and Tent5c^dcat/dcat^* (orange) mice. n=3 biological replicates per condition. **(d)** Representative micrographs of stage VII/VIII tubule cross sections from *Tent5c^null/null^* and *Tent5c^dcat/dcat^* mice compared to *Tent5c^wt/wt^*. INSL3 immunostaining in green; SYCP immunostaining (yellow) marks spermatocytes; PNA labeling (red) marks spermatid acrosomes, and DAPI labeling (grey) mark nuclei. Se, Sertoli cells; L, Leydig cells; ES, Elongated Spermatids. Scale, 20 μm. n=4-7 biological replicates per condition. **(e)** Box plot showing the quantification of the INSL3 immunostaining per Leydig cells from testis cross sections of *Tent5c^null/null^* and *Tent5c^dcat/dcat^* mice compared to *Tent5c^wt/wt^*. Data are expressed relative to *Tent5c^wt/wt^* mice. Each dot represents one mouse. The crosses display the means; the lines show the medians; the boxes indicate the first and third quartiles and the bars indicate the 10^th^ and 90^th^ percentiles. Ordinary one-way analysis of variance (ANOVA), Tukey’s mutilple comparison test. **p<0.01. n=2 biological replicates per condition. **(f)** Poly(A) tail length density plot as in (a) for *Insl3 and Ptgds* transcripts from whole testis of *Tent5c^wt/wt^* (magenta) *and Tent5c^dcat/dcat^* (orange) busulfan-treated mice. n=2 biological replicates per condition.

*Insl3* transcripts and proteins are highly accumulated in Leydig cells, but their role in sustaining spermatogenesis after testis descent remains unclear ^34^. A closer inspection of our immunostained samples revealed a decrease in INSL3 accumulation in the Leydig cells of *Tent5c^dcat/dcat^* mice compared to wild-type, as recently shown by Brouze and colleagues ^5^(**Fig. 4d and e**). To understand if the decrease in INSL3 protein levels in Leydig cells was associated with changes in the *Insl3* poly(A) tail length in the absence of TENT5C activity, we enriched the testis of wild-type and *TENT5c^dcat/dcat^* mice with interstitial cells by depleting germ cells with busulfan (**Extended Data Fig. 4c**). Immunostaining confirmed a decrease in INSL3 protein accumulation in *Tent5c^dcat/dcat^* Leydig cells even after busulfan injection (**Extended Data Fig. 4d**). Sequencing of busulfan treated testis revealed *Insl3 and Ptgds* transcripts mostly accumulating with poly(A) tails ∼180-nt long in wild-type samples (**Fig. 4f**). In *Tent5c^dcat/dcat^* busulfan treated testis, *Insl3* and *Ptgds* transcripts showed gradual deadenylation down to ∼60 nts with a significant depletion of the ∼180-nt long tail population (**Fig. 4f**). Poly(A) tail shortening was associated with the downregulation of *Insl3* transcripts (**Extended Data Fig. 4e, Extended Data Table 1**). Together, these results indicate that TENT5C poly(A) polymerase is required to maintain or readenylate *Insl3* poly(A) tails up to ∼180-nt long in Leydig cells and ∼150-nt long in ES. In contrast to long tails in ES, *Insl3* transcripts with ∼180-nt long tails in Leydig cells were translationally active given the high accumulation of INSL3 protein in these cells (**Fig. 4d**). Thus, the decrease in INSL3 protein levels in the Leydig cells from *Tent5c* mutants could be explained by a depletion of the translationally active transcripts in the absence of TENT5C-mediated adenylation. As *Insl3* poly(A) tails get shorter, transcripts may be triggered to decay resulting in their downregulation.

### TENT5C activity extends the poly(A) tails of mRNAs encoding structural proteins required for flagellum morphogenesis

To further understand the physiological relevance of TENT5C polyadenylation for sperm morphogenesis, we revisited our list of putative TENT5C targets in RS and ES. In addition to *Insl3* and *Ptdgs*, *Odf1 and Smcp* were identified as two of the four transcripts showing a significant reduction in poly(A) tail length in both *Tent5c^dcat/dcat^* spermatids (this study) and Tent5c*^null/null^* whole testis ^5^(**Fig. 5a**). ODF1 is a core component of the outer dense fibers (ODFs), a cytoskeletal structure which runs externally to the axoneme and provides the flagella with elastic properties and resistance to shear forces. As a result, ODF1 depletion leads to altered ODF arrangement, sperm decapitation and sterility ^6^. SMCP localizes to the mitochondrial capsule and is thought to support their spiral arrangement in the sperm midpiece ^35^. Therefore, *Odf1* and *Smcp* are candidates to causally explain the headless phenotype observed in *Tent5c* mutant sperm.

**Figure 5.**
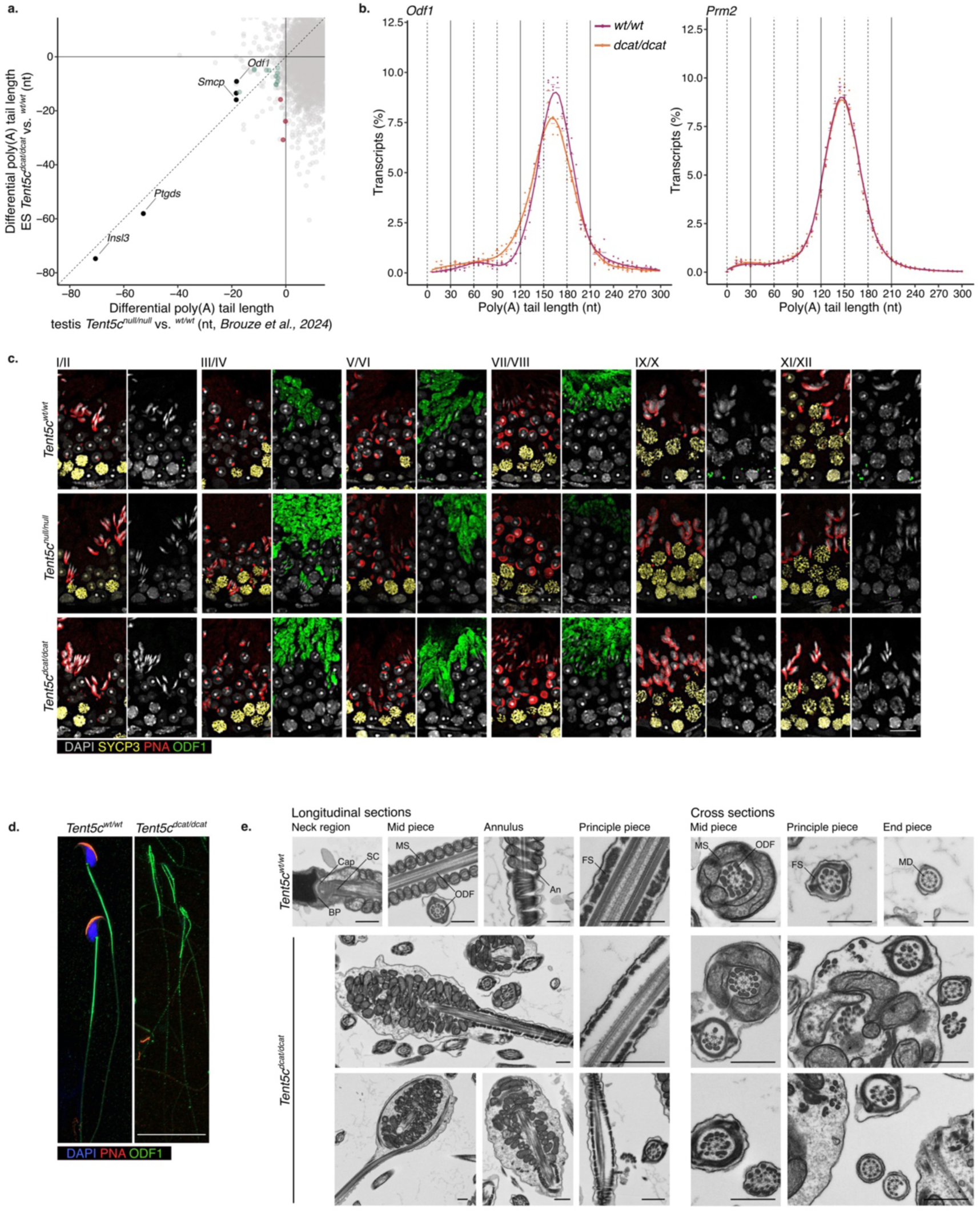
TENT5C activity extends the poly(A) tails of mRNAs encoding structural proteins required for flagellum morphogenesis. **(a)** Scatter plot comparing the differential poly(A) tail length of individual transcripts from the elongated spermatids (ES) of *Tent5c^wt/wt^* and *Tent5c^dcat/dcat^* mice (this study) and the differential poly(A) tail length of the corresponding transcripts from the whole testis of *Tent5c^wt/wt^* and *Tent5c^null/null^* mice ^5^. Each dot represents an individual transcript. Transcripts significantly decreasing in poly(A) tail length in elongated spermatids (ES) or whole testis are shown in purple and green respectively. The transcripts showing significance in both studies are shown in black and labelled by their gene symbol. nt: nucleotide. n=3 biological replicates per condition. **(b)** Poly(A) tail length density plot of *Odf1* and *Prm2* in elongated spermatid (ES) from *Tent5c^wt/wt^* (magenta) *and Tent5c^dcat/dcat^* (orange) mice. Dots indicate values for individual biological replicates. The bars indicate the relative mean percentage of transcripts for each poly(A) tail length. The local polynomial regression fitting is shown as a solid line for each condition. n=3 biological replicates per condition. **(c)** Representative micrographs of stage I/II to XI/XII tubule cross sections from *Tent5c^null/null^* and *Tent5c^dcat/dcat^* mice compared to *Tent5c^wt/wt^*. ODF1 immunostaining in green; SYCP immunostaining (yellow) marks spermatocytes; PNA labeling (red) marks spermatid acrosomes, and DAPI labeling (grey) mark nuclei. Scale, 20 μm. n=2-4 biological replicates per condition. **(d)** Representative micrographs of sperm from the cauda epididymis of *Tent5c^dcat/dcat^* mice compared to *Tent5c^wt/wt^*. ODF1 immunostaining in green; PNA labeling (red) marks sperm acrosomes, and DAPI labeling (blue) mark nuclei. Scale, 20 μm. n=2 biological replicates per condition. **(e)** Representative transmission electron micrographs of sperm from the caude epididymis of *Tent5c^dcat/dcat^* mice compared to *Tent5c^wt/wt^*. BP: basal plate; Cap: capitulum; SC: striated colums; MS: mitochondrial sheath; ODF: outer dense fiber; An: annulus; FS: fibrous sheath; MD: microtubule doublets. Scale, 600 nm.

As for the transition proteins and protamine mRNAs, the post-transcriptional regulation of *Smcp* and *Odf1* transcripts is characterized by changes in their poly(A) tail processing ^1,3^. While in RS, *Smcp* and *Odf1* transcripts accumulate with tails ∼150-nt long and are associated with the non-polysomal fraction, in ES, they enter the polysomal fraction with tails ∼150-nt long which get gradually shortened ^1,3^. Consistently, our sequencing methodology recapitulated the changes in *Smcp* and *Odf1* poly(A) tail length observed in the transition from RS to ES in wild-type mice (**Fig. 2c**). To understand the impact of TENT5C on the poly(A) processing of *Smcp* and *Odf1*, we examined their poly(A) tail profile in ES from *Tent5c^dcat/dcat^* animals.

*Smcp* and *Odf1* showed a decrease in transcripts with ∼150-nt poly(A) tails together with an increase in ∼60-nt long tails in ES expressing the catalytically mutant TENT5C when compared to wild-type ES (**Fig. 5b, Extended Data Fig. 5**). The shortening of *Smcp* and *Odf1* poly(A) tails observed in the absence of TENT5C activity is comparable to the one observed for the same transcripts during spermiogenesis progression, where a mild accumulation of ∼60-nts long tails reflects the translational activation of long-tailed mRNAs ^1,3^ (**Fig. 2c and d**). In contrast, *Prm2*, which shares a similar poly(A) profile to *Smcp* and *Odf1* in the transition from RS to ES, showed no poly(A) profile alteration in *Tent5c^dcat/dcat^* ES (**Fig. 5b**). Consistently, no changes in TNP2 protein expression were observed in TENT5C-depleted spermatids ^4^. Thus, the poly(A) polymerase activity of TENT5C is specifically required to maintain *Smcp* and *Odf1* transcripts with ∼150-nt poly(A) tails.

To interrogate possible changes in the ODF1 protein accumulation during spermiogenesis, we immunostained testis sections from wild-type and *Tent5c* mutants. In wild-type mice, ODF1 staining was first detected in step 15 spermatids and remained elevated until spermiation (**Fig. 5c**). The staining appeared diffuse across the cytoplasm with a concentrated signal in a single locus. Throughout spermatogenesis, ODF1 granules were observed at the base of seminiferous tubules, likely reflecting the elimination of ODF1 in residual bodies, as previously described in rats ^36^(**Fig. 5c**). In *Tent5c^null/null^* and *Tent5c^dcat/dcat^* mice, we observed consistent defects in ODF1 protein accumulation and distribution. In spermatids step 15, the cytoplasmic intensity of ODF1 signal was higher than in wild-type mice, with an irregular distribution and multiple loci (**Fig. 5c**). The frequency of ODF1-positive granules was much reduced, with only occasional isolated single granules (**Fig. 5c**). To determine whether the aberrant ODF1 accumulation was preserved in mature sperm, we immunostained sperm isolated from the cauda epididymis. In wild-type mice, ODF1 localized at the base of the sperm head, distributed uniformly along the midpiece to gradually decrease in intensity along the principal piece (**Fig. 5d**). In *Tent5c^dcat/dcat^* mice, ODF1 signal was heterogenous along malformed midpieces, clustering at sites where the flagella fused to form hairpins (**Fig. 5d**). Aberrant loci of high intensity were detected along the principal piece, down to the distal end of the flagella (**Fig. 5d**). These results show that TENT5C poly(A) polymerase activity is required to control the localized distribution of ODF1 protein in ES and sperm, suggesting that ODF1 dysregulation may contribute to the morphological defects associated with *Tent5c* mutants.

A previous study proposed the disassembly of the connecting piece linking the sperm head and flagellum as a main mechanism explaining the phenotype of TENT5C-depleted sperm^4^. To further investigate the effects of TENT5C deficiency —and ODF1 dysregulation— on sperm ultrastructure, sperm pellets extracted from the cauda epididymis were prepared for transmission electron microscopy. Sperm from wild-type mice showed a typical organization of the head-tail coupling apparatus, with discernable basal plate, capitulum and segmented columns (**Fig. 5e**). Only headless spermatozoa were captured in *Tent5c* deficient mice. The midpiece of wild-type spermatozoa showed a typical helical coiled sheath of mitochondria organized around a regular assembly of ODF tightly linked to the nine outer doublet microtubules of a unique axoneme (**Fig. 5e**). In contrast, TENT5C-deficient flagella presented disturbed structural organization of the mitochondrial sheath. Mitochondria formed multilayered aggregates, and multiple axonemal structures, often incomplete and asymmetric, were observed within the same cytoplasm (**Fig. 5e**). Longitudinal sections revealed looping of irregular axonemal structures around disorganized mitochondria. Additionally, a mix of mitochondria and fibrous sheath suggested abnormal annulus formation in the mutant cells (**Fig. 5e**). Aberrations in the principal piece of mutant sperm included disorganized or absent microtubule doublets, along with missing or misaligned ODFs (**Fig. 5e**). In several cross-sections, the ODFs were displaced to the inner side of the microtubule doublet, not in direct contact with the fibrous sheath (**Fig. 5e**). Longitudinal sections occasionally revealed gaps between the axoneme and fibrous sheath leading to protrusions along the principal piece (**Fig. 5e**). These defects closely resembled those characterized in *Odf1* mutant mice ^6^. Taken together, our results suggest that the poly(A) tail polymerase activity of TENT5C is required to regulate the spatial translation of ODF1 during spermiogenesis; a role essential to maintain sperm morphological integrity to support fertility.

## DISCUSSION

Transition proteins, protamines, *Smcp*, and *Odf1* mRNAs are transcribed in RS, but they only become translationally active as spermatids elongate ^1–3^. This temporal separation between transcription and translation suggests that long-term mRNA stabilization is essential for proper spermiogenesis ^8,9^. Although poly(A) tails are thought to contribute to RNA storage, observations are mostly limited to a few transcripts ^10–12^. Using global and quantitative approaches, we found that from spermatocytes to round spermatids, most transcripts gain longer poly(A) tails. The factors mediating the establishment of the ∼150-nt long poly(A) tails in RS remain unclear. The testis-specific poly(A) polymerase tPAP/PAPOLB, which predominantly accumulates in pachytene spermatocytes and RS, extends the poly(A) tails of *Ldhc*, *Odc1*, and specific transcription factor mRNAs in RS ^10,12^. Although tPAP activity is essential to support spermatids development, the extent of its activity across the transcriptome is unknown ^10^. TENT5C is unlikely to drive the polyadenylation of the RS transcriptome as it is expressed later in ES and targets only a few transcripts. The emergence of longer poly(A) tails in RS may also result from the *de novo* synthesis of mRNAs with long tails stabilized at ∼150 nts. Understanding how the RS transcriptome is stabilized with ∼150-nt long tails is required to gain insights into mRNA metabolism during spermiogenesis progression.

During spermatid elongation, the translational activation of previously studied transcripts correlates with poly(A) tail shortening ^1–3^. Our study shows that this association applies to the whole transcriptome during spermiogenesis. We observed that the emergence of ∼60-nt tails is a robust indication of active translation. However, the intensity of the translational activation is proportional to the abundance of transcripts retained with ∼150-nt tails. Consistently, previous work has shown that most *Smcp* and *Prm2* transcripts associated with high polysomal fractions have tails ∼150-nt long ^3^. Thus, translational activation occurs prior to deadenylation and does not require poly(A) tail shortening. Instead, the shortening from ∼150- to ∼60-nt long tails during spermiogenesis appears to be a byproduct of active translation. In this way, comprehensive poly(A) tail profiles can be used as specific and sensitive resources to infer the temporal dynamics of translation during spermiogenesis. For example, *Prm2* transcripts encode for the last of the transition proteins and protamines to appear in late spermatids ^37^. As such, *Prm2* transcripts primarily accumulate with poly(A) tails ∼150-nt long in ES, in contrast to the transition proteins and *Prm1* mRNAs where the fraction of transcripts with ∼150-nt long tails is much reduced.

Proteins like PABPC1 and the Y box proteins control the translation of mRNAs in spermatids ^38–41^. In the cytoplasm, PABPC1s prevent mRNA deadenylation by coating the poly(A) tail, with each PAPBC1 covering ∼27 nts ^42^. Although PABPC1s can favor translation, in excess they compete with the translational activation of spermatid mRNAs, ultimately leading to sterilityc^38^. Thus, in spermatids, 5 to 6 PABPC1s may bind and protect the transcript poly(A) tails from deadenylation and regulate the translational fate of mRNAs. Y-Box protein 2 (YBX2) and Y-Box protein 4 (YBX4) repress the translation of transcripts with long poly(A) tails in spermatids ^39–41^. Upon YBX2 depletion, the premature entry of *Prm1* and *Smcp* into the polysomal fraction results in spermiogenesis arrest ^39,40^. Similarly, maintaining high levels of YBX4 prevents the normal translational activation of transcripts such as *Tnp2* and outer dense fiber of sperm tails 2 (*Odf2*) ^41^. The translational repression by YBX2 and YBX4 may prevent the premature deadenylation of mRNAs in RS, securing their storage for protein synthesis in ES.

How the poly(A) tail polymerase activity of TENT5C may control mRNA metabolism in spermatids remains unclear ^4,5^. Here, we show that TENT5C activity is required to maintain the length of *Insl3* and *Ptgds* poly(A) tails at ∼150 nts in ES, but the redistribution of *Insl3* tails to ∼60-nt in *Tent5c^dcat/dcat^* ES was not associated with *Insl3* translational activation. These observations support that tail shortening is not a prerequisite for active translation. In Leydig cells, *Insl3* and *Ptgds* accumulate with ∼180-nt long tails, ∼27 nts longer than in spermatids, indicating that they are regulated by a different ribonucleoprotein (RNP) complex with one additional PABP ^42^. Unlike the RNP to which *Insl3* is associated in spermatids, the Leydig complex renders *Insl3* translationally active as shown by the presence of INSL3 protein in the interstitial compartment. As observed in spermatids, the maintenance of *Insl3* and *Ptgds* poly(A) tails above 150 nts depends primarily on TENT5C. In the absence of TENT5C poly(A) polymerase activity, both the protein and the mRNA levels of *Insl3* decrease, supporting a model where TENT5C activity is required to maintain translationally active transcripts. In the absence of TENT5C, the shortening of *Insl3* tails may contribute to transcript destabilization and decay.

INSL3 and PTGDS are both required for testis descent; *Insl3*- and *Ptgds*-deficient animals are cryptorchid and hemi-cryptorchid, respectively ^43–45^. Since *Tent5c* mutants do not show obvious aberrations in testis position, TENT5C does not appear to play a critical role during testicular development. The role of INSL3 in sustaining adult spermatogenesis is unclear ^34^. *Insl3* deficient animals are sterile due to cryptorchidism, but their fertility can be recovered if the testes are artificially descended, albeit with very low efficiency ^46^. Relaxin/insulin-like family peptide receptor 2 (RXFP2) is the cognate receptor for INSL3. Global *Rxfp2* knock-out mice are cryptorchids as expected; however, when *Rxfp2* is specifically removed from the germline, males sire normal-size litters ^47,48^. These observations indicate that INSL3 depletion does not directly impact spermatids. Given the morphological abnormalities identified in *Tent5c^dcat/dcat^* sperm, we also examined TENT5C targets encoding for structural proteins.

ODF1 and SMCP are proteins required for the morphological integrity of the flagella ODFs and mitochondrial sheath, respectively ^6,49^. ODF1-deficient animals show headless spermatozoa, a phenotype resembling that of TENT5C mutants ^6^. We identified *Odf1* and *Smcp* as putative targets of TENT5C poly(A) polymerase activity in ES. Consistently, both transcripts show a poly(A) tail shortening of similar amplitude in global testis samples of *Tent5c* null mice ^5^. Although the changes in poly(A) tail in mutant animals were modest, a reduction in *Odf1* and *Smcp1* poly(A) tail length of comparable magnitude was observed upon translational activation during spermiogenesis. Therefore, we deem these transcripts to be high-confidence TENT5C targets and propose that the shortening of *Odf1* and *Smcp* poly(A) tails in the absence of TENT5C activity impacts their translation.

ODF1 protein is not detected until spermatid stage 16; from this stage onwards, the protein appears concentrated in a single foci within the cytoplasm until elongated spermatids are released from the seminiferous epithelium. Positive ODF1 granules are then observed close to the basement membrane, presumably within residual bodies phagocytized by Sertoli cells ^36^. In TENT5C-deficient spermatids, ODF1 is mostly detectable by spermatids stage 16; however, the initial burst of translation is more pronounced and diffused throughout the cytoplasm. Thus, TENT5C poly(A) polymerase activity is essential to control the local translation of ODF1. ODF1 mislocalization could impair its direct interaction with factors necessary for the flagellum architecture. Consistently, the comprehensive characterization of sperm morphological defects by EM showed that the *Tent5c* mutant phenotype recapitulates abnormalities found in mutants for proteins interacting with ODF1. ODF1 binds the ODFs to the axoneme and the mitochondrial sheath by interacting with sperm associated antigen 4 (SPAG4) and kinesin light chain 3 (KLC3), respectively ^50,51^. Depletion of SPAG4 results in the partial disassembly of the axonemal complex as observed in animals encoding a catalytically dead TENT5C ^52^. Overexpression of a KLC3 mutant unable to interact with ODF1 results in midpiece abnormalities ^51^.

In elongated spermatids, ODF1 is abundant at the head-tail coupling apparatus and at the ODFs surrounding the axoneme, two structures derived from the centrioles ^36,53,54^. In addition, ODF1 is found at the manchette, a transient microtubular structure that guides the organization of the cytoskeletal elements required for spermatid elongation ^53^. Like ODF1, TENT5C also localizes to the manchette ^4^ and directly interacts with the centriolar protein PLK4 ^55,56^. These observations support a model where TENT5C poly(A) polymerase activity promotes the translation of *Odf1* transcripts next to the centriole and along the manchette, ensuring ODF1 proper distribution as the cytoplasm and flagella elongate. In the absence of TENT5C activity, the loading of ODF1 is more erratic as observed in the flagella of mature sperm, leading to a range of morphological abnormalities such as the cytoplasmic enlargement of the midpiece.

The translational control of mRNAs at centrioles is a conserved regulatory mechanism across animals ^57^. In the Drosophila embryo, Oo18 RNA-binding protein supports the local synthesis of Pericentrin-like protein at the centrosomes by promoting polyadenylation ^58^. Similarly, we speculate that the poly(A) polymerase activity of TENT5C anchors *Odf1* transcripts to the centriole and locally elongates their poly(A) tails to sustain translation. In the absence of TENT5C activity, *Odf1* transcripts diffuse away from the centriole and engage in translation, leading to their poly(A) tail shortening and the accumulation of ODF1 protein across the cytoplasm. As the translationally active pool of *Odf1* transcripts drops, the overall accumulation of ODF1 decreases as spermiogenesis progresses, contributing to the largely diminished presence of ODF1 foci close to the basement membrane of *Tent5c* mutants.

In summary, our study comprehensively documents the transcriptome-wide changes in poly(A) tail during spermiogenesis. Consistent with previous observations made on a few transcripts, a global increase in transcript poly(A) tail length in the transition from spermatocytes to spermatids is followed by a gradual shortening of poly(A) tails during spermatid elongation ^1–3,10–12^. While most mRNAs exhibit similar trends, each transcript displays unique variations in poly(A) profile reflective of specific mRNA stability and translation efficiency during spermiogenesis. In addition, our study shows the critical contribution of the poly(A) polymerase activity of TENT5C to support fertility by targeting a few transcripts in a cell-type- and organelle-specific manner. The deadenylation of *Odf1* affects the spatio-temporal distribution of the ODF1 protein required for flagellum morphogenesis, explaining how TENT5C polymerase activity supports sperm development and fertility. Given that the etiology of male infertility cannot always be determined by genetic variations or differences in RNA accumulation, our findings show that poly(A) tail length profiling can be used as a powerful tool to screen changes in transcript metabolism causative of sperm abnormalities.

## METHODS

### Mouse models

The following mouse strains were maintained to generate experimental animals: C57BL/6J (The Jackson Laboratory, IMSR_JAX:000664) for backcross and fertility test; tg(CMV-Cre) (B6.C-Tg(CMV-cre)1Cgn/J) (The Jackson Laboratory, IMSR_JAX:006054) for global *null* conversion of *floxed* alleles ^1^; *Tent5c* C-terminal-GFP knock-in, *Tent5c^gfp/gfp^* (C57BL/6-Tent5c<tm1(EGFP)Adki>) ^2^; loxP-flanked *Tent5c, Tent5c^fl/fl^* (B6.Cg-Tent5c<TM1MMO>) used to generate global *Tent5c* knockout mice; *Tent5c^null/null^* (B6.Cg-Tent5c<tm1.1Mmo>) and catalytically dead TENT5C, *Tent5c^dcat/dcat^* (C57BL/6-Tent5c<TM2ADKI>) ^2^. Mice were socially housed at the animal facility of the National Institute of Environmental Health Sciences and maintained on a 12 hours/12 hours light/dark cycle at 50 % ± 15 % humidity and 20.5 °C – 23.9 °C with free access to food and water. Animal studies were performed in accordance with the Guide for the Care and Use of Laboratory Animals from the National Institute of Health. Animal protocols were approved by the Institutional Animal Care and Use Committee.

### Generation of transgenic mice

The *Tent5c* conditional null (“flox”) locus was generated by inserting loxP sites in intron 1-2 and the 3’UTR, flanking the single coding exon of *Tent5c*. Two separate plasmid donors were used for each loxP sites. The 5’ loxP site donor plasmid consisted of a 400 bp 5’ homology arm (chr3:100,473,608-100,474,007 mm10), 430 bp 3’ homology arm (chr3:100,473,173-100,473,602 mm10), and a genetic payload of 42 bp that included the loxP site and unique restriction enzyme sites (SphI, NsiI, and NdeI). The 3’ loxP site donor plasmid consisted of a 461 bp 5’ homology arm (chr3:100,471,978-100,472,438 mm10), 379 bp 3’ homology arm (chr3:100,471,596-100,471,974 mm10), and a genetic payload of 43 bp that included the loxP site and unique restriction enzyme sites (KpnI, PsiI, and SspI). Gene targeting was done in B6129F1 embryonic stem cells (G4; 129S6/SvEvTac x C57BL/6Ncr). ES cells were lipofected with a 6:6:1:1 molar ratio of donor plasmids and two Cas9-Puro/sgRNA(5’ loxP TACTCTTGGTCGCAGCCGTG <u>NGG</u> and 3’ loxP TAGCACATGGGATCTGAGCT <u>NGG</u>) delivery plasmid (**pSpCas9(BB)-2A-Puro (PX459) V2.0**), a gift from Feng Zhang (Addgene plasmid #62988) ^3^. Following transfection, the cells were exposed to 48-hours of puromycin selection (0.9 µg/mL) followed by standard clonal expansion and screening. Clones were screened with 5’ and 3’ PCR screens external to the four homology arms to identify clones homozygous for the proper integration of both loxP sites: Tent5c 5’Scr (Fwd: CTTTTGGAAGTTTACCTGCCCG, Rev: CACAGAACCACATCTCTCACCA) and Tent5c 3’Scr (Fwd: GGATTTTGAGGAAGCCTTTGACC, Rev: GTTTCCTATTGACAATCACCGCC). Screening amplicons from targeted clones were fully sequenced to confirm homozygous insertion of each loxP site. Homozygous flox *Tent5c* ES cells were microinjected into albino B6J blastocyst for chimeric founder generation. The *Tent5c flox* allele was re-screened in the chimera F1 offspring. The line was then crossed to C57BL/6J wildtype mice to establish and expand the colony. To confirm proper Cre-mediated recombination and generate global *Tent5c* knock-out mice *Tent5c^null/null^*, the *Tent5c flox* mouse line was crossed to tg(CMV-Cre) mice from Jackson Labs (JAX stock #006054) ^1^. The closest likely Cas9 off-target location was a 3 bp mismatch located 65 Mbp away in intergenic sequence, so no attempts were made to screen for linked Cas9-mediated mutations.

### Genotyping

For routine colony maintenance, genotyping was performed by Transnetyx (Memphis, TN, USA) using automated DNA isolation from ear punches, followed by real-time PCR. The genotype of experimental animals was confirmed using conventional PCR from tail DNA. Briefly, 5 mm tail snip was digested at 56 °C in 200 µL proteinase K buffer until completely lysed. The DNA was precipitated with the addition of 200 µL of AL Buffer (DNeasy Blood & Tissue kit, Qiagen) and 200 µL of ethanol. The mixture was loaded on a DNeasy Mini spin column, and the DNA captured in the filter was sequentially washed in AW1 and AW2 buffers prior elution in 100 µL AE Buffer. DNA concentrations were determined by Nanodrop and adjust to 10 ng/µl for polymerase chain reaction. PCR were performed using ReadyMix Taq PCR Reaction Mix (Sigma-Aldrich, P4600). 20 µL enzymatic mix was prepared according to the manufacturer’s protocol and 20 ng of DNA added to the reaction for amplification. The primer sequences, number of cycles, annealing temperature and extension time specific to each amplification are provided in **Extended Data Table 2**. For each reaction, water was used as a non-template control. When specified, the PCR products were digested with Pfe1 or BamH1 restriction enzymes (NEB) according to the manufacturer’s instructions. Amplicons were run together with nucleic acid ladders on a 1.4% agarose gel with ethidium bromide. Imaging was processed using the Amersham ImageQuant800 imaging system (Cytiva).

### Testicular Single-Cell Suspensions

Testis isolated from 8-12 weeks male mice were decapsulated and collected in 25 mL of a freshly prepared Enriched Krebs-Ringer bicarbonate (EKRB) medium maintained at room temperature (120.1 mM NaCl, 4.8 mM KCl, 25.2 mM NaHCO_3_, 1.2 mM KH_2_PO_4_, 1.2 mM MgSO_4_.7H_2_O, 11 mM Glucose, 1.3 mM CaCl_2_, 1X Pen/Strep (Sigma, P0781), 1X Essential AA (Sigma, M71450), pH 7.4). Seminiferous tubules were dissociated from the interstitial tissue by Type XI Collagenase digestion (0.5 mg/ml, Sigma, C7657) in a 34 °C water bath for a minimum of 10 minutes with periodic tube inversion until tubules appeared dispersed. Tubules were then allowed to sediment for 2 minutes at room temperature and the interstitial cells were removed with the supernatant. Tubules were washed once with 25 mL of plain EKRB to remove residual collagenase and interstitial cells. The seminiferous tubules were further digested in 10 mL of EKRB supplemented with Trypsin-EDTA (0.005 %, Gibco, 25200-056) in a 34 °C water bath for 15 minutes. After incubation, tubules were dissociated by pipetting 10–15 times with a serological pipet until a single-cell suspension was obtained. Trypsin activity was then stopped quickly by the addition of 1 mL of FBS (10 % final concentration) and the solution of cells was centrifuged 500 g for 5 minutes at RT to clear out digestion media. The cell pellet was resuspended in 10 mL EKRB-10 % FBS and cell suspension was passed through a 0.45 uM nylon cell strainer to remove cell clumps. A 20 µL aliquot of cell suspension was removed for cell counting (Cellometer Auto T4 system, Nexcelom Bioscience). Cells were centrifuged at 500 g for 5 minutes at RT and resuspend with EKRB-10 % FBS at a concentration of 1 million cells /mL. Aliquots of 4 million cells were set aside to control for no staining, Hoechst 33342 staining only and PI staining only. The rest of the cell suspension was sequentially stained with Hoechst 33342 (3.2 ug/mL, Invitrogen, H3570) at 34 °C for 20 minutes, and PI (1 ug/mL, Invitrogen, P3566) at 34 °C for an additional 10 minutes. Stained cells were immediately centrifuged at 500g for 5 minutes at 4 °C. A suspension of 13 million cells/mL in staining media was prepared and maintained at 4 °C until FACS sorting.

### Flow Cytometric Analysis and Cell Sorting

Cells were sorted according to their characteristic Hoechst fluorescence and light scattering using a BD FACSAriaII cell sorter (Becton Dickinson Biosciences) equipped with a 70-micron nozzle and FACSDiVa software. Initially, gates were set on a side-scatter (SSC-H vs SSWC-W) followed by a forward scatter (FSC-H vs FSC-W) dot plot to isolate single cells. These cells were projected onto a forward scatter (FSC-A) versus side scatter (SSC-A) to identify the principal population of single cells free of debris. Dead cells were excluded using PI (Ex: 561; Em: 585) on a PI (PE-A) histogram. One hundred thousand viable cells were recorded for analysis. Viable cells stained with Hoechst excited using a UV (Ex: 355) laser were initially analyzed on a UV-A (605/40; 595 LP) versus UV-B (450/50) dot plot. Three distinct populations of cells were gated: Gate I for leptotene/zygotene spermatocytes (L/Z spermatocytes), Gate II for P/D spermatocytes and Gate III for haploid spermatids. The Gate III population was subsequently examined on a FSC-A versus SSC-A dot plot to identify the Gate IIIa and IIIb populations corresponding to RS and ES respectively. Cell sorting was performed capturing the populations from the Gate I, II, IIIa and b. In some experiments, the gates were projected onto a FITC (Ex: 488; 525/50) histogram to capture GFP positive cells. Cells were sorted into EKRB-10% FBS pre-coated 1.5 ml Eppendorf tubes. Collected cells were smeared on positively charged glass slides (Superfrostplus, Thermo Scientific) for histological analyses. Dry pellets were snap frozen after centrifugation at 500 g for 25 minutes at 4 °C for molecular analyses.

### RNA extraction

Frozen testis or germ cell pellets were homogenized in 1 mL of TRIzol and allowed to reach room temperature. Five minutes later, 200 µL of chloroform-isoamyl alcohol were added. Then, the mixture was vigorously shaken, incubated for 2 minutes at room temperature, and centrifuged at 12,000 g for 15 minutes at 4 °C. The aqueous fraction was moved to a new tube, and RNA was recovered by adding 400 µL isopropanol and 0.5 µL of GlycoBlue to facilitate the visualization of the RNA pellets. After 15 minutes, the samples were centrifuged at 12,000 g for 10 minutes at 4 °C. The RNA pellet was washed with 75 % ethanol, then centrifuged at 7,500 g for 5 minutes at 4 °C. After ethanol removal, total RNA was resuspended in nuclease-free water and quantified using Qubit (Qubit™ RNA HS Assay Kit, Thermofisher, Q32852). RNA integrity was confirmed on the bioanalyzer system Agilent 2100 (Agilent Technologies) prior to further analysis.

### Poly(A) tail purification

For Direct RNA sequencing libraries, mRNAs were purified from total RNA using Dynabeads™ Oligo(dT)_25_ beads (ThermoFisher). 20 µL of oligo(dT) beads per sample were washed with 40 µL of binding buffer (20 mM Tris-HCl, pH 7.5, 1.0 M LiCl, and 2 mM EDTA) and resuspended in 40 µL of binding buffer. 1-4 µg of total RNA per sample were diluted to 40 µL in nuclease-free water and mixed with 40 µL of binding buffer, followed by denaturation at 65°C for 2 minutes. The denatured RNAs were mixed with 40 µL of washed oligo(dT) beads and incubated at room temperature for 5 minutes. Beads were collected on a magnetic rack and washed twice with wash buffer (10 mM Tris-HCl, pH 7.5, 0.15 M LiCl, and 1 mM EDTA). After the final wash, poly(A)-tailed mRNAs were eluted in 10 µL of elution buffer (10 mM Tris-HCl, pH 7.5) by incubating the beads at 80°C for 2 minutes.

### Library Multiplexing

For multiplexing of poly(A) tail libraries from different samples, we followed the same barcoded-tagged oligonucleotide strategy described by Smith and collaborators ^4^. Briefly, four unique barcode sequences (BC1–BC4) were incorporated in the splint-ligation oligonucleotides; oligonucleotide sequences are listed in **Extended Data Table 3**. To create barcoded oligonucleotide duplexes, partially reverse complementary oligonucleotides (final concentration of 1.4 µM each) were incubated together at 94°C for 5 minutes in annealing buffer (0.01 M Tris-HCl, pH 7.5, 0.05 M NaCl) and gradually cooled to room temperature (-0.1 °C/second). These barcoded oligonucleotide duplexes were used in the initial ligation step following the Oxford Nanopore Technologies Direct RNA sequencing protocol (ONT, SQK-RNA002). This barcoding strategy was used to prepare the following libraries: The P/D spermatocyte library multiplexes 3 mRNA extracts (BC 1-3), each representative of a pool of mRNAs from the P/D spermatocytes of 3 independent wild-type mice. The RS library multiplexes 3 mRNA extracts (BC 1-3), each representative of a pool of mRNAs from the RS of 3 independent wild-type mice. The busulfan libraries, replicate 1 and 2, are each multiplexing 4 mRNA extracts (BC 1-4) from 1 mouse per condition (*Tent5c^wt/wt^* and *Tent5c^dcat/dcat^*, vehicle or busulfan injected). Other libraries were designed to detect poly(A) tail length and terminal modifications. For this we followed a split-ligation-based strategy previously described ^5,6^. A barcoded-tagged duplex oligonucleotides mix was prepared by mixing different annealed barcoded oligonucleotide, with each oligonucleotide duplex designed to capture RNA terminal molecules by identifying associated barcode sequences. Ten sets of duplex annealed oligos were made by mixing complementary oligonucleotides (**Extended Data Table 3**) and incubating them at 94 °C for 5 minutes, followed by cooling to room temperature in the annealing buffer (0.01 M Tris-HCl, pH 7.5, 0.05 M NaCl). Then, a 100 nM oligonucleotide mix was prepared according to the ratios of A, U, and G capturing oligos listed in **Extended Data Table 3**. BC1 (Oligo 1 in the BC mix) was used for identifying non-modified poly(A) tails and BC2 (Oligo 2 in the BC mix) was used for identifying terminal mono-uridylation. BC3 (Oligos 3–6 in the BC mix) and BC4 (Oligos 7–10 in the BC mix) were used for identifying oligo-uridylation and terminal guanylation, respectively. This barcoding strategy was used to prepare the RS and ES libraries from mutant mice and their corresponding wild type. 2 to 3 libraries per genotype and cell type were generated. Each library is built of a pool of mRNAs isolated from the spermatids of 3 independent mice. Each library includes four distinct barcodes (BC 1-4) to detect 3’-end residues as described above.

### Direct RNA sequencing

Direct RNA sequencing libraries were prepared according to the ONT standard protocol (Direct RNA Sequencing Kit, SQK-RNA002), with modifications to incorporate barcoded oligonucleotides. To capture poly(A) tail length and terminal modifications, a splint ligation method was used to ligate barcoded oligonucleotides to the 3′ ends of transcripts. For this, a split-ligation reaction was set up in a 15 µL volume by adding 9 µL of poly(A)-enriched mRNA, 3 µL of ligation buffer (6 % PEG8000 in 10× T4 DNA ligation buffer, NEB), 1.5 µL of T4 DNA ligase (NEB), 1 µL of 0.1 ng/µL RNA standards mix ^7^, and 1.5 µL of annealed barcoded oligonucleotides (100 nM) for library multiplexing or barcoded-oligonucleotides duplex mix (100 nM) for poly(A) tail terminal modifications (**Extended Data Table 3**). The reaction was incubated for 10 minutes at room temperature to anneal barcoded oligonucleotides for multiplexing, or 60 minutes at room temperature to enhance ligation efficiency when using barcoded oligonucleotides for terminal modifications. To maximize sequencing throughput, the ligated RNA was reverse-transcribed into an RNA-cDNA hybrid according to the manufacturer’s instructions (Thermo Fisher Scientific). Briefly, 15 µL of split-ligated RNA was mixed with 9 µL of nuclease-free water, 2 µL of dNTPs (10 mM), 8 µL of 5X first-strand buffer, 4 µL of 100 mM DTT, and 2 µL of SuperScript II reverse transcriptase (Thermo Fisher Scientific). The reaction mix was incubated at 50 °C for 50 minutes, followed by 10 minutes at 70 °C and cooling to 4 °C in a thermocycler. The resulting RNA-cDNA hybrid was purified using 72 µL of RNAClean XP beads (Beckman Coulter, A63987) according to the manufacturer’s instructions, washed with 150 µL of 70% ethanol, and eluted in 20 µL of nuclease-free water on a magnetic stand. For multiplexing, distinct barcodes were eluted together after 5 minutes at room temperature in 20 µL of nuclease-free water. The RNA sequencing adapters containing motor protein (RMX, supplied in the Direct RNA Sequencing Kit SQK-RNA002) were ligated to the ends of RNA-cDNA hybrids. Briefly, 8 µL of ligation buffer (6% PEG in 10× T4 DNA ligation buffer, NEB), 6 µL of RMX-RNA adapter, 3 µL of nuclease-free water, and 3 µL of T4 DNA ligase (NEB) were mixed with 20 µL of purified RNA-cDNA hybrid in a final reaction volume of 40 µL. The ligation reaction was incubated for 15 minutes at room temperature. Adapter-ligated RNA-cDNA hybrids were purified using 40 µL of AMPure beads (Beckman Coulter), washed twice with 150 µL of wash buffer (WSB from SQK-RNA002), and eluted from beads in 21 µL of elution buffer (EB from SQK-RNA002) after a 10-minute incubation at room temperature. The libraries were prepared for loading into ONT flow cells (R9) using the Flow Cell Priming Kit (EXP-FLP002) according to the manufacturer’s instructions. Briefly, 21 µL of adapter-ligated libraries were mixed with 37.5 µL of RNA running buffer (RRB from SQK-RNA002) and 17.5 µL of nuclease-free water. The R9 flow cell was primed by adding 800 µL of the FLT/FB mix (30 µL FLT in 1 mL FB tube, from EXP-FLP002) into the priming port, followed by an additional 200 µL after 5 minutes of incubation. Finally, 75 µL of the library was loaded through the spot-on port and sequenced on a MinION or GridION instrument (ONT).

### Fertility test

To test for fertility, each male mice 8-12 weeks old were housed with two virgin C57BL/6J/B6 female mice 6-8 weeks old for 5 days. Females were monitored for pregnancies and isolated for delivery. The number of pups per litter was recorded on the day of birth. For each male mouse tested the number of pups per litter was average between the 2 females.

### Busulfan treatment

To significantly decrease the proportion of germ cells in mouse testis and enrich for somatic cells, a freshly prepared busulfan solution (2.5 mg/mL in a 1:1 mixture of dimethyl sulfoxide and distilled water), or the corresponding vehicle, were injected intraperitoneally to *Tent5c^wt/wt^* or *Tent5c^dcat/dcat^* adult male mice in a single dose of 20 mg/kg (0.2 mL for a 25 g mouse). Testis were harvested 4 weeks after injection to allow for an adequate depletion of germ cells before histological and molecular analysis ^8^.

### Sperm isolation and count

Both epididymes were collected in a 20 mm round bottom glass dish containing 500 µL of 34 °C EKRB buffer (120.1 mM NaCl, 4.8 mM KCl, 25.2 mM NaHCO3, 1.2 mM KH2PO4, 1.2 mM MgSO4.7H2O, 11 mM Glucose, 1.3 mM CaCl2, 1X Pen/Strep (Sigma, P0781), 1X Essential AA (Sigma, M71450), pH 7.4) supplemented with 10 % FBS. The cauda epididymis was isolated and punctured four times with a 26-gauge needle. The dish was incubated at 34 °C for 10 minutes to allow the spermatozoa to swim out. The epididymal tissue was removed and the sperm suspension were transferred into a clean low binding Eppendorf tube with an additional 500 µL of EKRB buffer for a total volume of 1 mL. Sperm suspension was centrifuged at 500 g for 5 minutes with the brakes set to minimum to avoid cell damage. Sperm pellet was then resuspended in 500 µL PBS 1X. Droplets corresponding to 20 µL sperm suspension were smeared on positively charged glass slides (Superfrostplus; Thermo Scientific) for histological analyses. The original sperm suspension was diluted 10 times in PBS 1X and loaded onto a 0.100 mm deep Malassez hemocytometer. The number of flagella, used as a proxy to quantify germ cell production, was counted manually under a brightfield microscope (Axio, Zeiss).

### Tissue and smear processing

For histological analysis testes and epididymes were fixed in 4 % paraformaldehyde solution at 4°C overnight and paraffin embedded. 5 μm sections were mounted on positively charged glass slides (Superfrostplus, Thermo Scientific), deparaffinized, and rehydrated prior staining or immunohistochemistry. Spermatozoa and sorted cells were smeared on positively charged glass slides (Superfrostplus, Thermo Scientific), allowed to dry over-night, fixed in 4% paraformaldehyde solution for 15 minutes and rehydrated.

### Conventional histological staining

For H&E staining, tissue sections and spermatozoa smears were stained in Mayer’s hematoxylin for 4 minutes and washed for 5 minutes in running water to remove excess dye and allow bluing. Testis and epididymis sections were then stained in eosin solution for 7 seconds and immediately transferred to 70 % ethanol. Spermatozoa smears were left in contact to eosin for 2 minutes to obtain a better contrast of the flagellum. For Periodic Acid-Schiff (PAS) staining, rehydrated testes sections were oxidized in 0.5% periodic acid solution for 5 minutes and rinsed 3 times in distilled water. Sections were then incubated for 15 minutes in Schiff’s reagent, washed with distilled water, counterstain in Mayer’s hematoxylin for 1 minute and quickly transferred to running water. Sections were then dehydrated with rising ethanol concentrations, transferred into xylene and permanently mounted in a xylene-based mounting medium. Brightfield images were captured on a Zeiss AxioObserver Z1 microscope (Carl Zeiss Inc). Transmitted light was collected with a Zeiss AxioCam IC color camera. The Zeiss Zen Blue software was used for image processing.

### Immunofluorescence

Rehydrated testis sections, spermatozoa and sorted cell smears were treated for antigen retrieval in 250 mL of boiling antigen unmasking solution (Vector, H3300) for 15 minutes, allowed to cool down for 20 minutes and washed in PBS 1X for 5 minutes. Cell membranes were permeabilized in a 0.4 % triton X-100 solution for 10 minutes before incubation in a blocking solution (PBS 1X, 0.4 % triton X-100, 1 % BSA, 5 % normal donkey serum) for 1 hour at room temperature. Slides were then incubated overnight at 4 °C with primary antibodies diluted in PBS 1X-0.4 % triton X-100 (INSL3, 1/200, ThermoFisher, PA5-55921; ODF1, 1/200, Abcam, ab197029) or no primary antibody (negative controls). To detect cell death, sections were further stained using the In Situ Cell Death Detection Kit Roche, #11684817910) and incubated 1 hour at 37 °C with TUNEL reaction mixture (50 μL of enzyme solution mixed into 450 µL Label solution just prior to use) following manufacturer’s instructions. Slides were then washed 3 times in PBS 1X-0.4 % triton X-100 and incubated for 1 hour at room temperature with secondary antibodies (1/1000), Rhodamine to label Peanut agglutinin in the acrosomes (PNA, 1/1000, Vector Laboratories, #RL-1072), and the Alexa Fluor® 647 Anti-SYCP antibody (1/200, Abcam, #ab205847) to identify meiotic cells. After 3 washes in PBS 1X, the TrueView Autofluorescence Quenching Kit (Vector Laboratories, #SP-8400-15) was used to reduce cell autofluorescence. DAPI mounting media was used to stain for nuclei (VECTASHIELD Vibrance® Antifade Mounting Medium with DAPI, Vector Laboratories, H-1800-10). Images were captured using the Zeiss LSM 780 UV confocal microscope. Tissue sections were scanned with a 2 µm interval Z stack followed by maximum intensity projection. Sperm images were captured with a Zeiss LSM 980 confocal with AiryScan for sperm. The laser power and gain settings were held constant for image comparison. The Zeiss Zen Blue software was used for selecting region of interest and image processing.

### Electron microscopy

Sperm were isolated from the cauda epididymis and pellets were fixed in Modified Karnovsky fixative at RT for approximately 2 hours. Samples were rinsed with 0.1 M phosphate buffer, post-fixed in 1% osmium tetroxide in phosphate buffer, rinsed in distilled water, dehydrated through a graded ethanol series (70 %, 75 %, 95 %, 100 %), and transitioned to acetone. The samples were then infiltrated with increasing concentrations of Polybed 812 resin (1:3 resin/acetone, 1:2, 1:1 resin/acetone to absolute Resin), embedded in absolute resin, and placed in an oven at 60 °C for 48 hours to polymerize. Once polymerized, blocks were trimmed and semithin cross-sections of the skin approximately 0.5 µm thick were cut, mounted on glass slides, and stained with 1 % toluidine blue O in 1 % sodium borate. The sections were examined by light microscopy to identify the areas of interest. Each block was trimmed to the area of interest and thin-sectioned at approximately 70 nm, placed on a 200 mesh copper grid, and stained with 3% aqueous uranyl acetate and Reynolds lead citrate. Digital images were captured with an AMT 16 megapixel camera attached to a JEOL JEM-1400+ transmission electron microscope operating at an accelerating voltage of 80 kV.

### Direct RNA sequencing analysis and poly(A) tail-length determination

Guppy (Oxford Nanopore Technologies) was used for read base calling. The reads were then aligned to the GENCODE vM17 (GRCm38.p6) genes using Minimap2 software v2.20-r1061, with the -x parameter set to map-ont ^9^. The length of the poly(A) tail was determined using Nanopolish ^10,11^. To run the Nanopolish poly(A) pipeline, reads were indexed with Nanopolish and mapped reads were sorted and indexed with SAMtools ^12^. Reads were demultiplexed using Deeplexicon ^4^. Only reads with poly(A) tail quality score PASS and confidence interval above 0.85 were selected for further analysis. The number of mapped reads per sample after quality check is reported in **Extended Data Table 4**. For further analysis, a minimal pre-filtering was performed to only consider the contigs with a minimum of 10 reads in at least 2 replicates in one of the conditions. The resulting files including transcript ID, poly(A) tail length, modification and replicate information for each read were used as an input for poly(A) tail analysis. The mean poly(A) tail length of each contig in each of the replicates was calculated and compared between conditions. A two-tailed t test was used to identify transcripts with poly(A) tails statistically shorter (-) or longer (+) between conditions (p-value ≤ 0.05). For each contig in each of the replicates, the number of reads with poly(A) tail length within 30 to 120 nts (reported as ∼60-nt poly(A) tails) and 120 to 210 nts (reported as ∼150-nt poly(A) tails) were quantified and expressed as a percentage relative to the total number of reads obtained from the same contig.

### Differential transcript accumulation analysis

The Bioconductor package DESeq2 was used for differential transcript accumulation analysis ^13^. Briefly, a count matrix was obtained from direct RNA sequencing using the DESeqDataSetFromMatrix function, and used as an input to build the DESeqDataSet file. The original DESeq functions were used for the standard differential expression analysis steps (normalization, dispersion estimation and fold change estimation) as described ^13^. Result tables were generated using the function results, which extracts log2 fold changes, Wald test p-values and adjusted p-values corrected for multiple testing using the Benjamini and Hochberg procedure. Result tables are shown in **Extended Data Table 1**. A significant threshold of q-value ≤ 0.05 was used for data interpretation. The plotPCA function was used to visualize the overall effect of experimental covariates on a PCA plot.

### Differential translation

Differential RNA translation was determined with Xtail v1.1.5 ^14^ using publicly available data reporting reads counts of ribosome profiling (Ribo-seq) and matched RNA sequencing (RNA-seq) libraries from adult mouse testis (ArrayExpress accession number: E-MTAB-7247) ^15^.

### Cell composition analysis

The count matrix generated by the DESeq2 workflow from direct RNA sequencing was used as an input for cell composition analysis using the Cell Population Mapping (CPM) deconvolution algorithm included in the scBio package ^16^. The abundance of testicular cell types for each sequenced replicates was inferred comparing expression profiles to a publicly available single-cell transcriptomic of mouse spermatogenesis used here as a reference (GSE accession number: GSE104556) ^17^.

### INSL3 immunofluorescence quantitation

Interstitial areas (6 to 10 per mouse) were isolated from images of testis sections stained for INSL3 and DAPI using the ROI function of the Zeiss Zen Blue software. These ROI images were then imported into FIJI (ImageJ version 1.53c). The green component of the RGB image corresponding to the INSL3 channel was extracted and converted to greyscale using the duplicate channel function. The interstitial compartment was then manually isolated from adjacent tubules with the freehand drawing tool followed by the clear outside function. The interstitial content was then converted to a binary mask to identify the INSL3 signal using the setAutoThreshold tool with the “Default dark no-reset” option. Watershed was applied to segment the binary image, and the particles were analyzed using the analyze particle command with the following options: size=0-Infinity circularity=0.00-1.00 show=Masks display exclude summarize add. The area of the particles corresponding to the INSL3 signal were expressed relative to the corresponding number of Leydig cells quantified manually from the DAPI channel.

### Statistical analysis

The statistical analysis details of each experiment can be found in the corresponding figure legend. To determine whether the data met the assumptions of the statistical tests, normal distribution and equal variance across samples were tested using a Shapiro-Wilk test and a Levene test, respectively. For non-parametric comparisons between more than two samples, a Krustal-Wallis one-way analysis of variance applying the Bonferroni correction for multiple testing was used. When the data met the normal distribution assumption, parametric tests were used. To compare two unpaired samples with equal variance a two-samples, two-tailed Student’s t-test was used. To compare more than two populations with equal variance, statistical analyses were performed using a one-way ANOVA followed by Tukey’s multiple comparison test.

## DATA AVAILABILITY

All the sequencing data are publicly available as of the date of publication and has been deposited at the Gene Expression Omnibus (GEO) ^18^ with the accession number GSE290698, GSE290699, GSE290700. Publicly available datasets used in this study: GENCODE vM17 (GRCm38.p6) (https://www.gencodegenes.org/mouse/release_M17.html); Single-cell transcriptomic of mouse spermatogenesis (GSE accession number: GSE104556) ^17^; Reads counts of ribosome profiling (Ribo-seq) and matched RNA sequencing (RNA-seq) libraries from adult mouse testis (ArrayExpress accession number: E-MTAB-7247) ^15^.

## ACKNOWLEDGEMENTS

We thank the members of the Morgan group for helpful discussions on the project. This research was supported by the Intramural Research Program of the NIH, National Institute of Environmental Health Sciences (grant ZIA ES103339 awarded to MM). This work was supported by the following NIEHS facilities: the flow cytometry center, the epigenomics and DNA sequencing core facility, the integrative bioinformatics support group, the gene editing and mouse model core facility, the animal resources section, the fluorescence microscopy and imaging center and the electron microscopy core laboratory. We thank Andrzej Dziembowski for sharing the *Tent5c^dcat^* and *Tent5c^gfp^* mouse lines generated by the Genome Engineering Facility part of the Inn-Mol-Cell infrastructure at the International Institute of Molecular and Cell Biology in Warsaw.

## AUTHORS CONTRIBUTIONS

MB performed most of the experiments and analyzed the data. AG contributed to the generation of the Nanopore direct RNA seq libraires. ACP contributed to the characterization of the Busulfan-treated testis. BP contributed to the analysis of the RNA sequencing libraries. MM and MB designed the study and wrote the manuscript. MM supervised the study.

## MATERIALS & CORRESPONDENCE

Original gel or microscopy pictures reported in this paper and any additional information required to reanalyze the data reported in this paper are available from the corresponding author upon request.

## EXTENDED DATA FIGURES AND TABLES

**Extended Data Figure 1.**
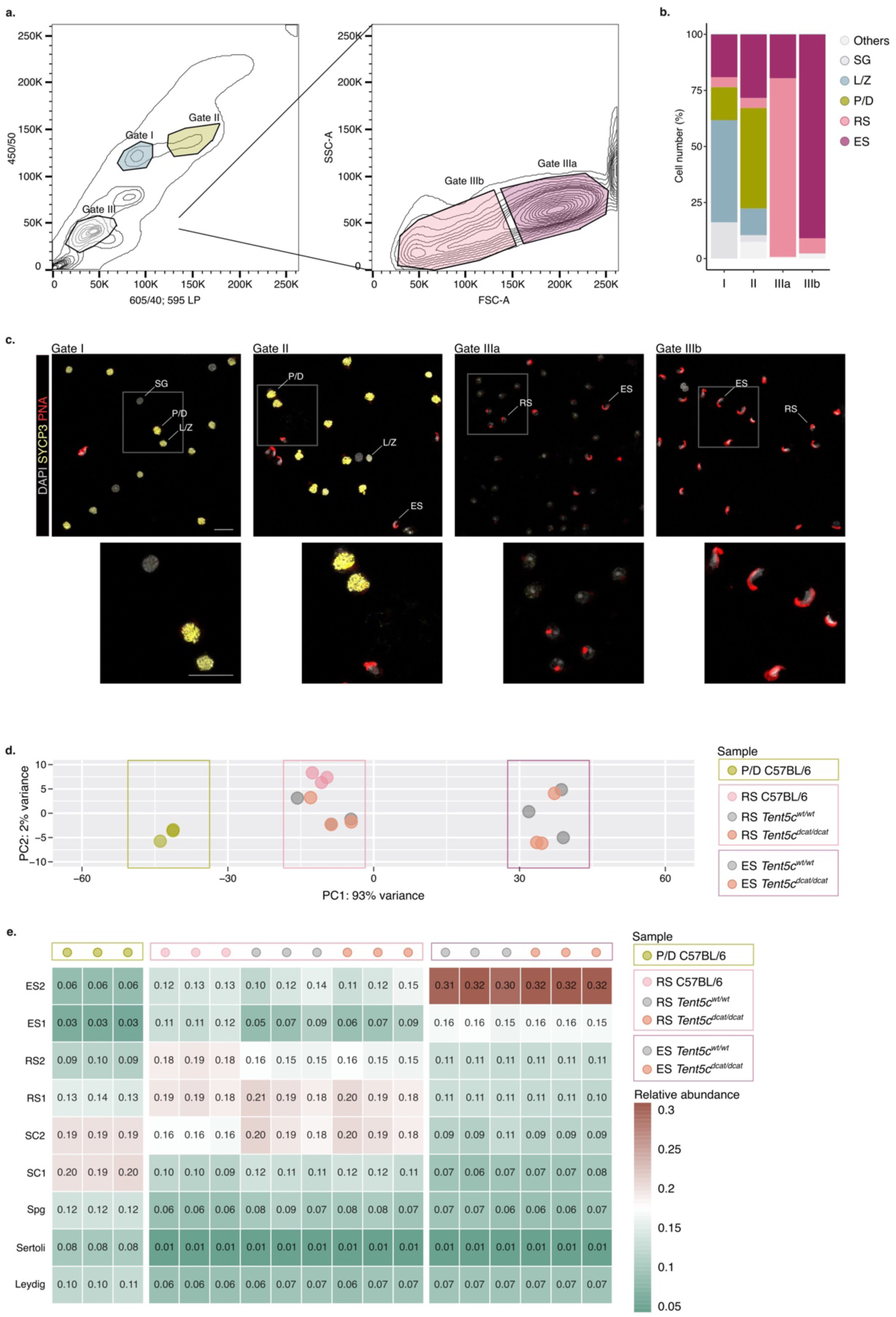
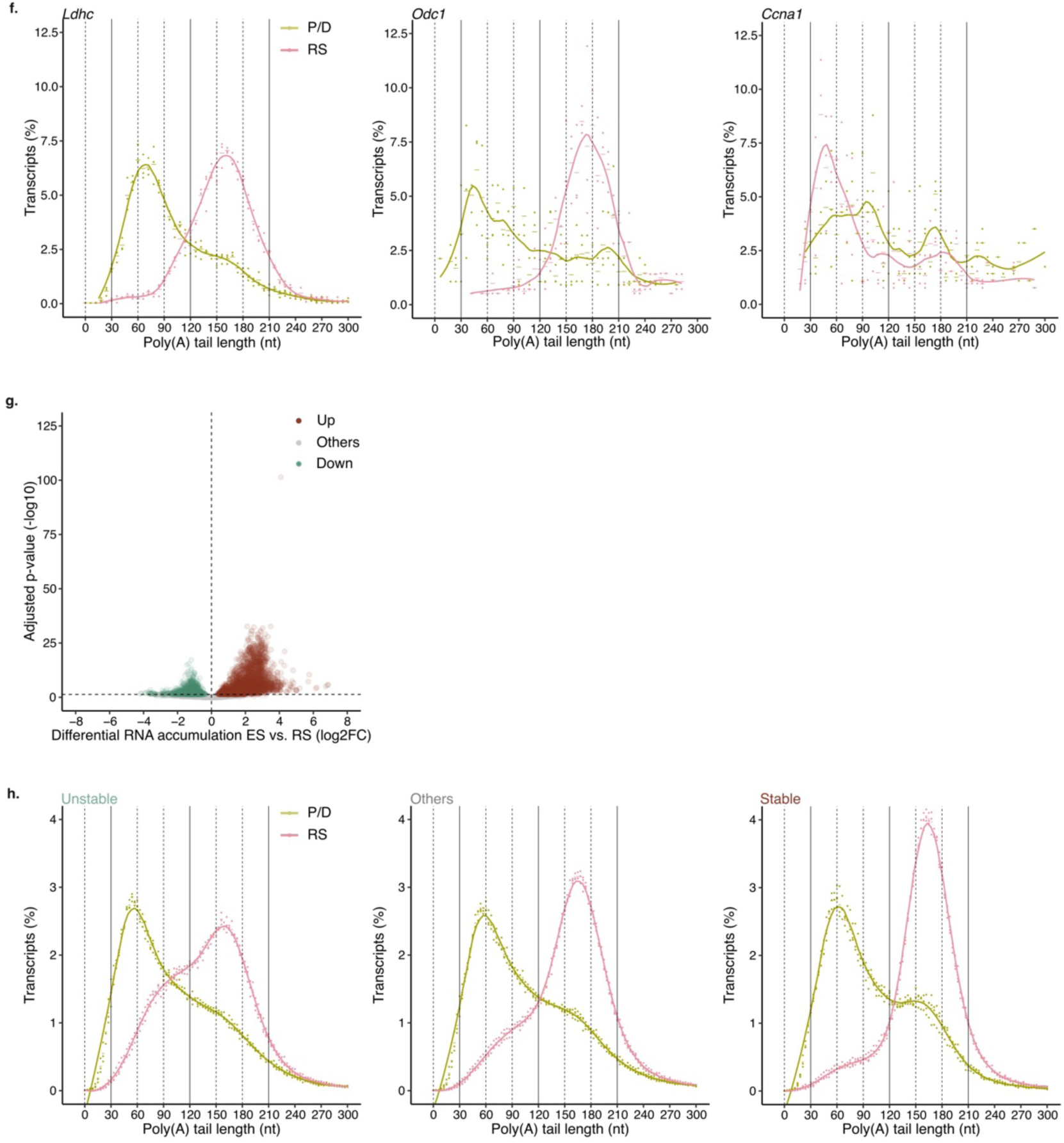
**(a)** Density plot representative of the gating strategy used in flow cytometry to isolate germ cell populations based on Hoechst fluorescence (left) and light scattering (right). **(b)** Bar plot showing the proportion of spermatogenic cells in the isolated populations from Gate I, **11,** Illa and lllb. SG, spermatogonia; UZ, leptotene/zygotene spermatocytes; P/D, pachytene/diplotene spermatocytes; RS, round spermatids; ES, elongated spermatids. **(c)** Representative micrographs of the germ cell populations isolated by flow cytometry in Gate I, II, Illa and lllb. SYCP immunostaining (yellow) marks leptotene/zygotene spermatocytes (UZ; punctate signal characteristic of unpaired synaptonemal complex) and pachytene/diplotene spermatocytes (P/D; elongated and intense signal indicating fully synapsed homologous chromosomes). **PNA** labeling (red) marks the acrosome apparatus of round spermatids (RS; punctate signal) and elongated spermatids (ES; elongated signal). DAPI labeling (grey) marks nuclei. Insets are highlighted in white. Scale, 20 µm. **(d)** Principal Component Analysis (PCA) representation of the covariates for all sequenced spermatogenic cell populations (pachytene/diplotene spermatocytes, PIO; round spermatids, RS and elongated spermatids, ES) isolated from C57BU6, Tent5cwVwt, and Tent5cdcaVdcat mice. n=3 biological replicates per condition. **(e)** Cell population mapping (CPM) showing the relative abundance of testicular cell types in all sequenced spermatogenic cell populations isolated from C57BU6, Tent5cwt/wt, and Tent5cdcaVdcat mice. n=3 biological replicates per condition. **(f)** Poly(A) tail length density plot for Ldhc, Odc1, and Ccna1 in pachytene/diplotene spermatocytes (PID, green) and round spermatids (RS, pink). Dots indicate values for individual biological replicates. The bars indicate the relative mean percentage of transcripts for each poly(A) tail length. The local polynomial regression fitting is shown as a solid line for each condition. nt: nucleotide. n=3 biological replicates per condition. **(g)** Volcano plot of differential RNA accumulation in elongated spermatids (ES) relative to round spermatids (RS). Upregulated (up), others, and downregulated transcripts (down) are indicated in brown, grey and green, respectively. Wald test corrected by Benjamini and Hochberg for multiple testing. Significance threshold q-val < 0.05. n=3 biological replicates per condition. (h) Poly(A) tail length density plots as in (f) for pachyteneldiplotene spermatocytes spermatocytes (P/D, green) and round spermatids (RS, pink) transcriptomes split by unstable (left), others (middle) and stable (right) transcripts. n=3 biological replicates per condition.

**Extended Data Figure 2.**
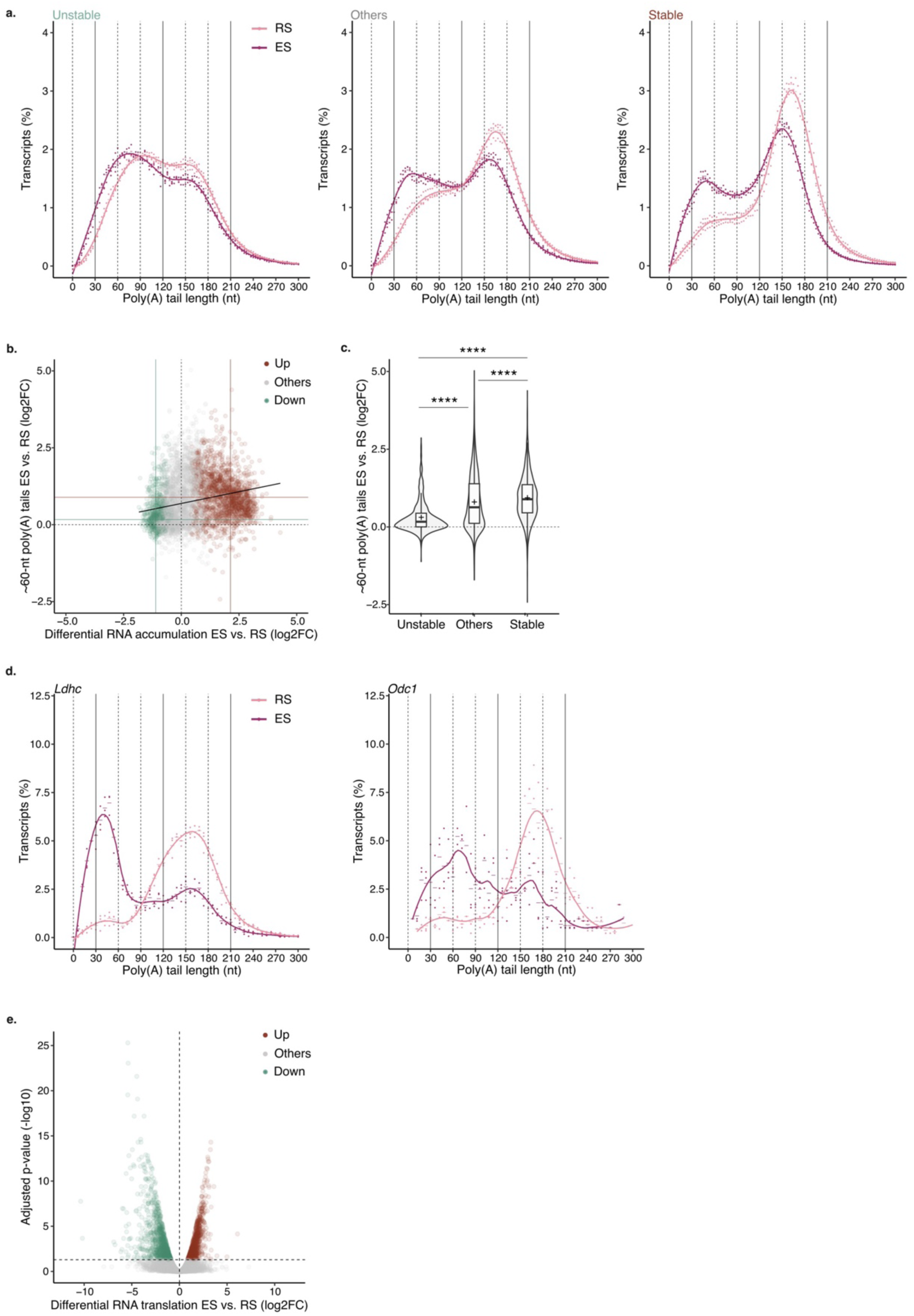
**(a)** Poly(A) tail length density plots for round spermatids (RS, pink) and elongated spermatids (ES, magenta) transcriptomes split by unstable (left), others (middle) and stable (right) transcripts. Dots indicate values for individual biological replicates. The bars indicate the relative mean percentage for each poly(A) tail length. The local polynomial regression fitting is show as a solid line for each condition. nt: nucleotide. **(b)** Scatter plot comparing fold change in transcript accumulation to the fold change in the proponion of reads per transcript with poly(A) tails ∼60-nt long between round (RS) and elongated spermatids (ES). Each dot represents an individual transcript. Upregulated (up), others, and downregulated transcripts (down) are indicated in brown, grey and green, respectively. Solid lines indicate the median of both variables for upregulated and downregulated mRNAs in brown and green, respectively. The linear fit is shown in black. **(c)** Violin plot showing the fold change in the proportion ot reads per transcript with poly(A) tails -60-nt long between round (RS) and elongated spermalids (ES) for transcripts grouped by stability. The width of violins show the density of individual transcripts. The overlaid box plots display means as crosses and medians as lines; the boxes indicate the first and third quartiles and the bars indicate the 10th and 90th percentiles. Krustal-Wallis test using Bonferroni correction for mutilple testing.**** P < 0.0001. **(d)** Poly(A) tail length density plot as in (a) for Ldhc and Odct in RS (pink) and ES (magenta). **(e)** Volcano plot for the differential RNA translation in elongated spermatids (ES) relative to round spermatids (RS). Transcripts with increased, unchanged or decreased level of translation are indicated in brown, grey and green respectively. Wald test corrected by Benjamini and Hochberg for multiple testing. Significance threshold q-val < 0.05.For all plots, n=3 biological replicates.

**Extended Data Figure 3.**
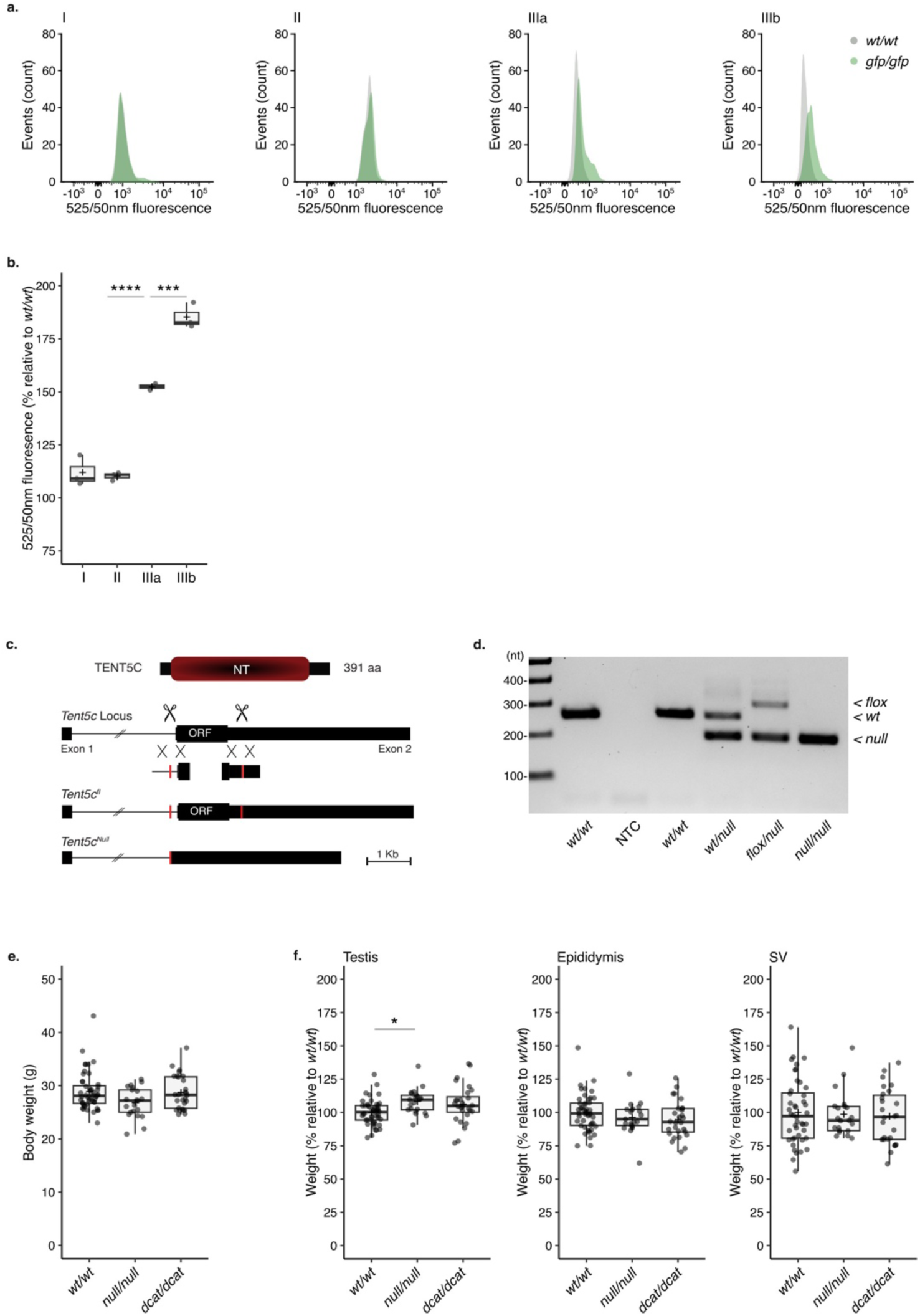
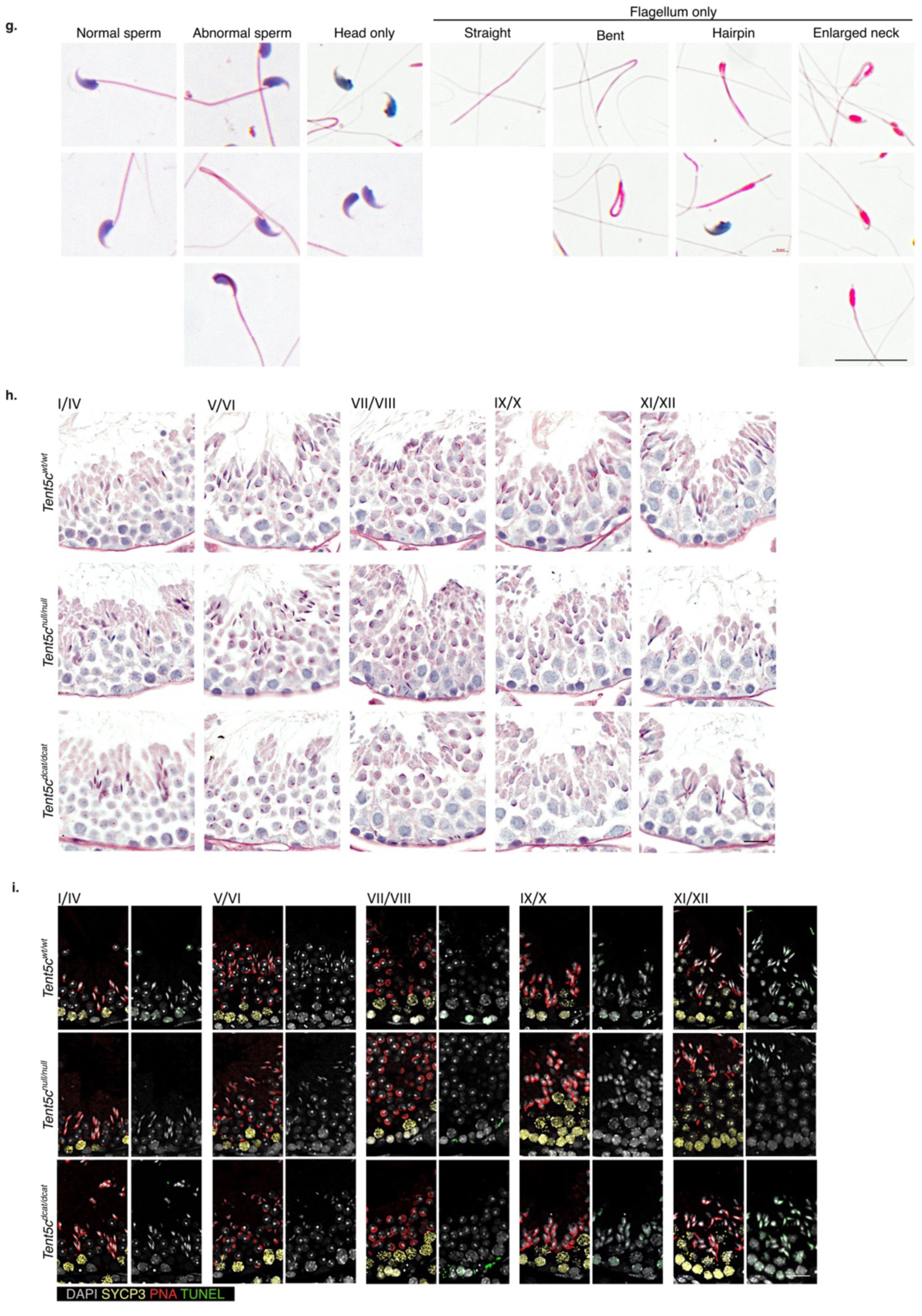
**(a)** Representative histograms of the intensity of 525/50 nm fluorescence in germ cell populations sorted from Tent5cwt/wt and Tent5cgfplgfp mice shown in grey and green respectively. n=3 biological replicates per condition. **(b)** Box plot showing the intensity of 525/50 nm fluorescence in germ cell populations sorted from Tent5cgfp/gfp mice. Data are expressed relative to Tent5cwtlwt mice. Each dot represents one mouse. The crosses display the means; the lines show the medians; the boxes indicate the first and third quartiles and the bars indicate the 10th and 90th percentiles. Ordinary one-way analysis of variance **(ANOVA),** Tukey’s mutilple comparison test. ***p<0.001, ****p<0.0001. n=3 biological replicates per condition. **(c)** Schematic of the targeting strategy used to develop the Tent5cflox and Tent5cnull alleles. (d) Representative gel images of the PCR amplification of the Tent5cwt, Tent5cflox and TentScnull alleles from DNA isolated from the tails of Tent5cwVwt, TentScwVnull, Tent5cflox/null and, TentScnull/null mice. The molecular marker is shown on the left. nt: nucleotide; NTC: non template control. (e) Box plot as in (b) showing the body weight of Tent5cwt/wt, Tent5cnulllnull and Tent5cdcaVdcat adult mice at sacrifice. Each dot represents one mouse. n=21-47 biological replicates per condition. **(f)** Box plots as in (b) showing testis, epididymis and seminal vesicle (VS) of Tent5cwt/wt, Tent5cnull/null and Tent5cdcatldcat adult mice. Organ weights are expressed relative to brain weight, and displayed as percentage relative to the average of the TentScwt/wt controls. Krustal-Wallis test using Bonferroni correction for mutilple testing. *p < 0.05. n=21-47 biological replicates per condition. **(g)** Representative micrographs showing the range of germ cell morphologies observed in the semen from the caudal epididymis of Tent5cwt/wt, Tent5cnulllnull and Tent5cdca/dcat mice atter Hematoxylin and Eosin (H&E) staining. Sperm heads (nuclei) show a blue staining and flagella shades of pink. Scale, 20 µm. (h) Representative micrographs showing stage I to XII tubule cross sections stained with Periodic Acid Schiff (PAS). Sections from Tent5cnulllnull and Tent5cdcat/dcat mice are compared to Tent5cwt/wt. The stage VIINIII tubule micrographs are also show in Figure 3E. Scale, 20 µm. n=21-47 biological replicates per condition. n=3 biological replicates per condition. (i) Representative micrographs as in (h) of stage I to XII tubule cross sections. SYCP immunostaining (yellow) marks spermatocytes; PNA labeling (red) marks spermatid acrosomes, TUNEL immunostaining (green) marks apoptotic cells and DAPI labeling (grey) marks nuclei. The stage VIINIII tubule micrographs are also show in Figure 3F. Scale, 20 µm. n=3 biological replicates per condition.

**Extended Data Figure 4.**
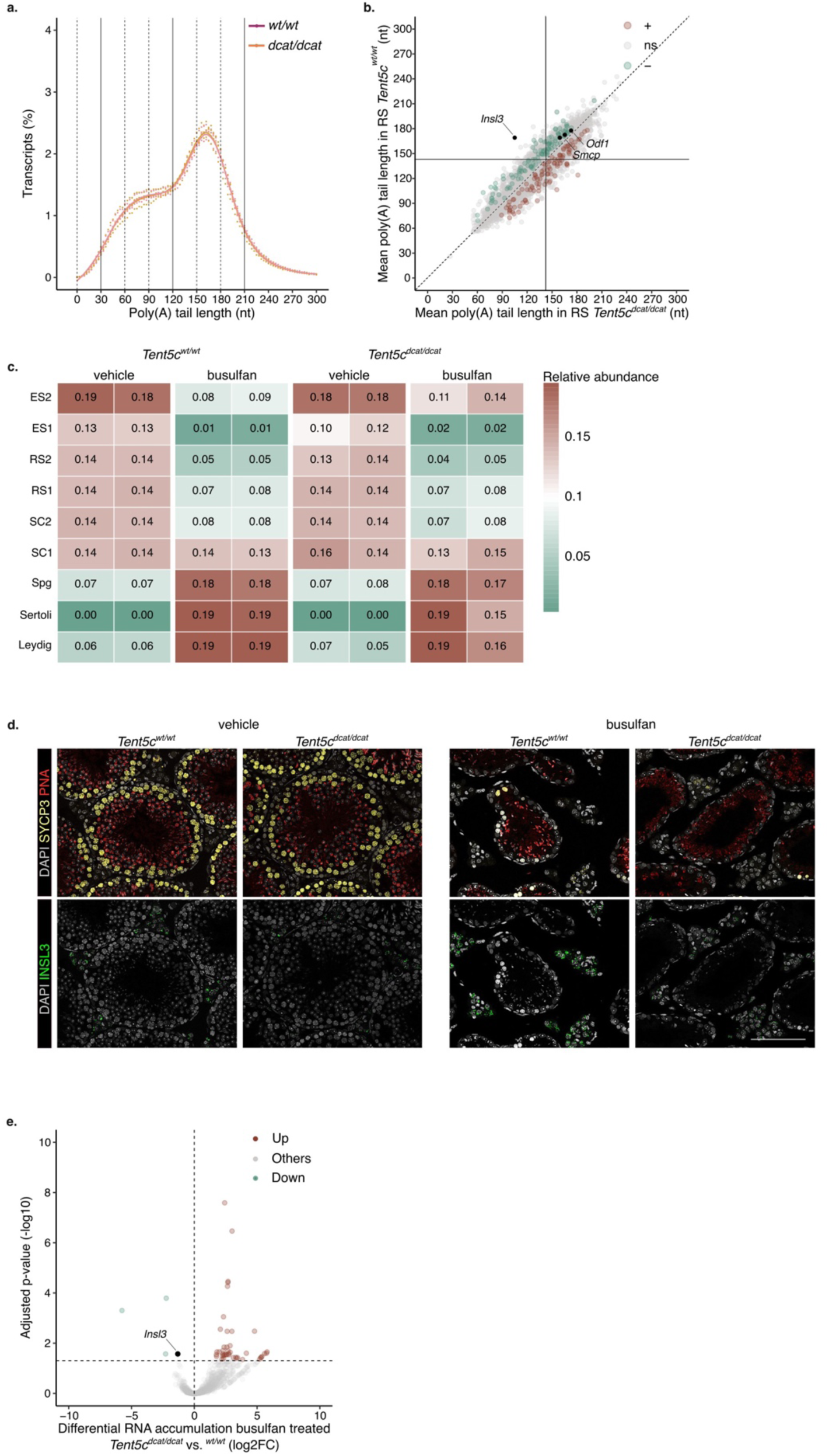
**(a)** Poly(A) tail length density plot of the round spermatid (RS) transcriptomes of Tent5cwt/wt (pink) and Tent5cdcat/ dcat (orange) mice. Dots indicate values for individual biological replicates. The bars indicate the relative mean percentage of transcripts for each poly(A) tail length. The local polynomial regression fitting is shown as a solid line for each condition. nt: nucleotide. n=3 biological replicates per condition. **(b)** Scatter plot comparing the mean poly(A) tail length of each transcript in round spermatid (RS) of Tent5cwt/wt and Tent5cdcat/dcat mice. Each dot represents an individual transcript. Transcripts significantly increasing(+) or decreasing(-) in poly(A) tail length between genotypes are shown in brown or green, respectively; Student’s t-test, two-tailed, significance threshold p < 0.05. ns: non significant. n=3 biological replicates per condition. (c) Cell population mapping (CPM) showing the relative abundance of testicular cell types in whole testis sequenced from Tent5cwt/wt and Tent5cdcatidcat mice 4 weeks after vehicle or busulfan (20 mg/kg) injection. n=2 biological replicates per condition. **(d)** Representative micrographs showing testis cross sections from Tent5cwt/wl and Tent5cdcat/dcat mice 4 weeks atter vehicle or busulfan (20 mg/kg) injection. INSL3 immunostaining in green: SYCP immunostaining (yellow) marks spermatocytes: PNA labeling (red) marks spermatid acrosomes, and DAPI labeling (grey) mark nuclei. Scale, 100 µm. n=2 biological replicates per condition. (e) Volcano plot of differential RNA accumulation in the busulfan treated testes of Tent5cdcat/dcat relative to Tent5cwt/wt mice. Upregulated (up), others, and downregulated transcripts (down) are indicated in brown, grey and green, respectively. lnsl3 transcript is indicated in black. Wald test corrected by Benjamini and Hochberg for multiple testing. Significance threshold q-val < 0.05. n=2 biological replicates per condition.

**Extended Data Figure 5.**
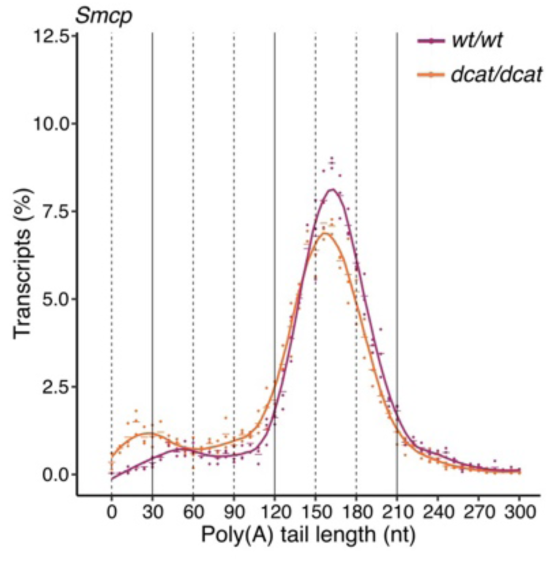
Poly(A) tail length density plot of Smcp in elongated spermatid (ES) from Tent5cwt/wt (magenta) and Tent5cdcat/dcat (orange) mice. Dots indicate values for individual biological replicates. The bars indicate the relative mean percentage of transcripts for each poly(A) tail length. The local polynomial regression fitting is shown as a solid line for each condition. nt: nucleotide. n=3 biological replicates per condition.

**Extended Data Table 2.**
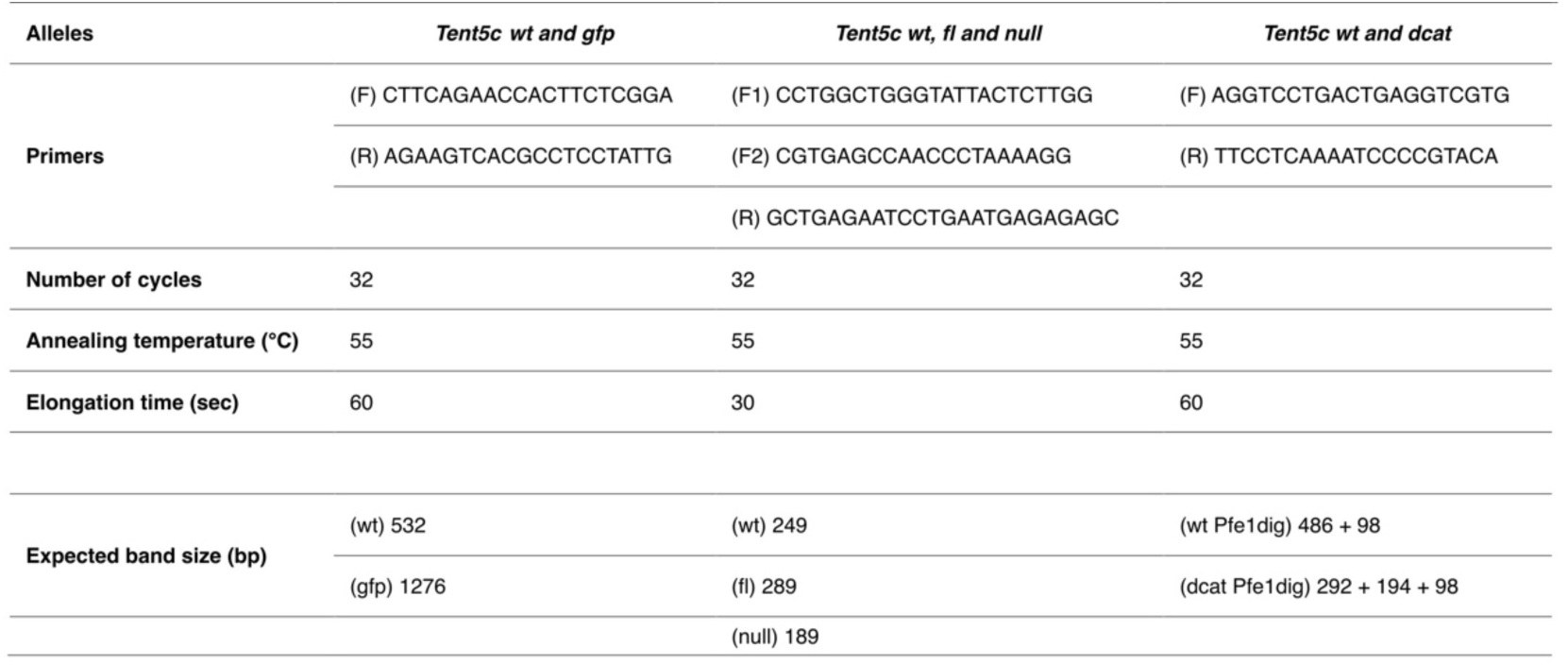
Genotyping primers and PCR conditions for allele amplification.

**Extended Data Table 3.**
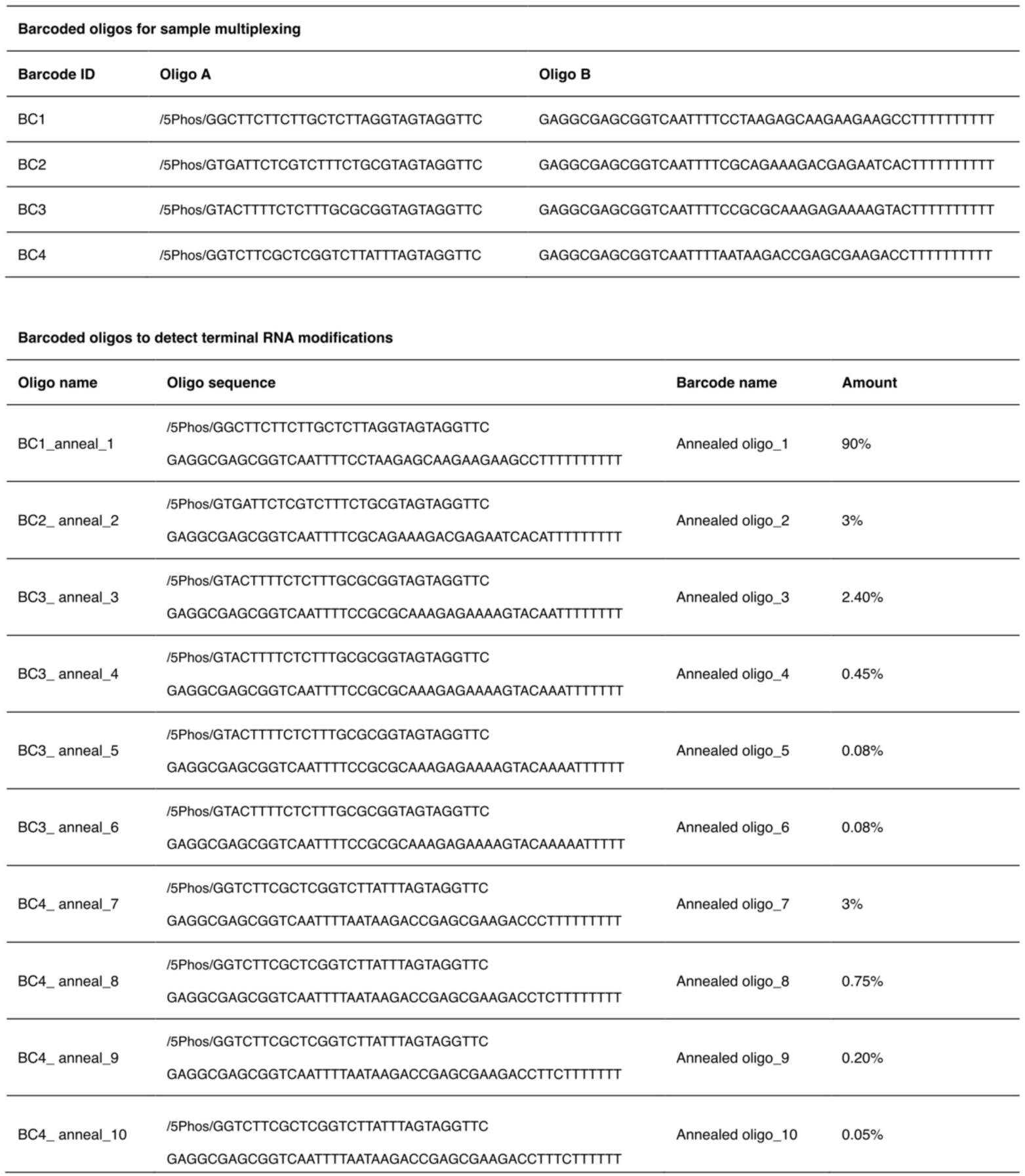
Splint oligos barcoded for Direct RNA Sequencing multiplexing.

**Extended Data Table 4.** Direct RNA Sequencing read count obtained after quality check (QC)

| Sample name | Read count after QC |
| --- | --- |
| pachytene_1 | 263890 |
| pachytene_2 | 330400 |
| pachytene_3 | 293310 |
| round_1 | 338725 |
| round_2 | 509157 |
| round_3 | 381851 |
| RS_TENT5c_WT_1 | 978889 |
| RS_TENT5c_WT_2 | 708550 |
| RS_TENT5c_WT_3 | 1455436 |
| RS_TENT5c_dCAT_1 | 784068 |
| RS_TENT5c_dCAT_2 | 647247 |
| RS_TENT5c_dCAT_3 | 1030092 |
| ES_TENT5c_WT_1 | 674834 |
| ES_TENT5c_WT_2 | 742072 |
| ES_TENT5c_WT_3 | 359278 |
| ES_TENT5c_dCAT_1 | 603182 |
| ES_TENT5c_dCAT_2 | 538070 |
| ES_TENT5c_dCAT_3 | 768682 |
| WT_veh_1 | 83105 |
| WT_veh_2 | 119739 |
| WT_bsf_1 | 101287 |
| WT_bsf_2 | 179537 |
| DCAT_veh_1 | 49267 |
| DCAT_veh_2 | 217305 |
| DCAT_bsf_1 | 96283 |
| DCAT_bsf_2 | 154287 |

